# Hijacking Methyl Reader Proteins for Nuclear-Specific Protein Degradation

**DOI:** 10.1101/2022.01.24.477424

**Authors:** Dhanusha A. Nalawansha, Ke Li, John Hines, Craig M. Crews

**Affiliations:** Department of Molecular, Cellular and Developmental Biology, Yale University, New Haven, Connecticut 06511, United States; Department of Chemistry, Yale University, New Haven, Connecticut 06511, United States; Department of Pharmacology, Yale University, New Haven, Connecticut 06511, United States

## Abstract

Targeted protein degradation (TPD) by PROTACs is a promising strategy to control disease-causing protein levels within the cell. While TPD is emerging as an innovative drug discovery paradigm, there are currently only a limited number of E3 ligase: ligand pairs that are employed to induce protein degradation. Herein, we report a novel approach to induce protein degradation by hijacking a methyl reader: E3 ligase complex. L3MBTL3 is a methyl lysine reader protein that binds to the Cul4^DCAF^^5^ E3 ligase complex and targets methylated proteins for proteasomal degradation. By co-opting this natural mechanism, we report the design and biological evaluation of L3MBTL3-recruiting PROTACs and demonstrate nuclear-specific degradation of FKBP12 and BRD2. We envision this as a generalizable approach to utilize other reader protein-associated E3 ligase complexes in PROTAC design to expand the E3 ligase toolbox and explore the full potential of TPD.

## Introduction

Targeted Protein Degradation (TPD) by PROteolysis TArgeting Chimeras (PROTACs) has emerged as an exciting area in both basic biological discovery as well as in drug development^1–3^. PROTACs are heterobifunctional molecules comprised of a target binding ligand connected to an E3 ligase recruiting ligand via an intervening linker^4–5^. PROTACs induce proximity-dependent ubiquitination and the subsequent proteasomal degradation of the target protein, hence eliminating all functions of the protein. Given their unique mode of action, PROTACs offer advantages over small molecule inhibitors^6–7^, e.g., catalytic activity and the ability to address non-enzymatic functions of proteins. Most PROTACs to date degrade target proteins by utilizing either VHL or cereblon as the recruited E3 ligase^8^. This focus on a limited number of E3 ligases is due to the paucity of liganded E3 ligases in the proteome to recruit via a small molecule-based PROTAC. Therefore, expanding the E3 ligase toolbox is an active area of research in TPD^9^. Since co-opting E3 ligases for PROTAC development could lead to tumor-specific, tissue-specific and compartment-specific protein degradation, it is important to explore novel E3 ligases with differential expression patterns and/or distinct functions in order to mitigate the limitations associated with current PROTACs, such as on-target toxicity and the emergence of resistance to CRBN- and VHL-based PROTACs^10–14^.

Recent studies have used chemoproteomic methods to identify covalent ligands for novel E3 ligases, and demonstrated their potential in PROTAC design^15^. In 2019, the Cravatt lab identified a covalent DCAF16 ligand by screening electrophilic scout fragments, and subsequently demonstrated nuclear-specific FKBP12 and BRD4 degradation by PROTACs structurally based on the covalent ligand^16^. More recently, the Cravatt lab has identified DCAF11 as another ligandable E3 ligase by screening electrophilic PROTACs derived from a more diversified scout library^17^. The Nomura lab has also identified covalent ligands for multiple E3 ligases such as RNF114, RNF4, KEAP1 and FEM1B, and demonstrated their applicability in TPD by targeting BRD4 for degradation^18–22^. While the use of chemoproteomic strategies to discover covalent E3 ligase ligands has expanded the repertoire of E3 ligases in PROTAC design, the development of innovative strategies to recruit novel E3 ligases with unique features for TPD is still warranted.

E3 ubiquitin ligases induce conditional protein ubiquitination by recognizing a specific degron in the target protein, such as phospho-degron, N- and C-degrons, hydroxyproline degrons and methyl degrons^23–24^. Among these, methyl degrons are one of the more poorly studied. The methyl degron was first described in 2012, where methylation-dependent ubiquitination and degradation of the tumor suppressor RORα was controlled by the Cul4^DCAF1^ E3 ligase^25^. While several studies showed that lysine methylation induces protein degradation, the mechanism was not clearly defined^26–29^. Fortunately, in 2018, two studies identified that the methyl-lysine reader protein, L3MBTL3 (Lethal(3)Malignant Brain Tumor-Like Protein 3), in complex with the Cul4^DCAF5^ E3 ligase, targets methylated proteins such as DNMT1, SOX2 and E2F1 for proteasomal degradation (Figure 1A)^30–31^. Based on this evidence as well as the success seen with co-opting the hydroxyproline-dependent degradation by VHL, we explored whether hijacking a methyl-lysine reader proteins to recruit the E3 ligase complex to the target protein and explore their utility and implications in TPD (Figure 1B).

**Figure 1.**
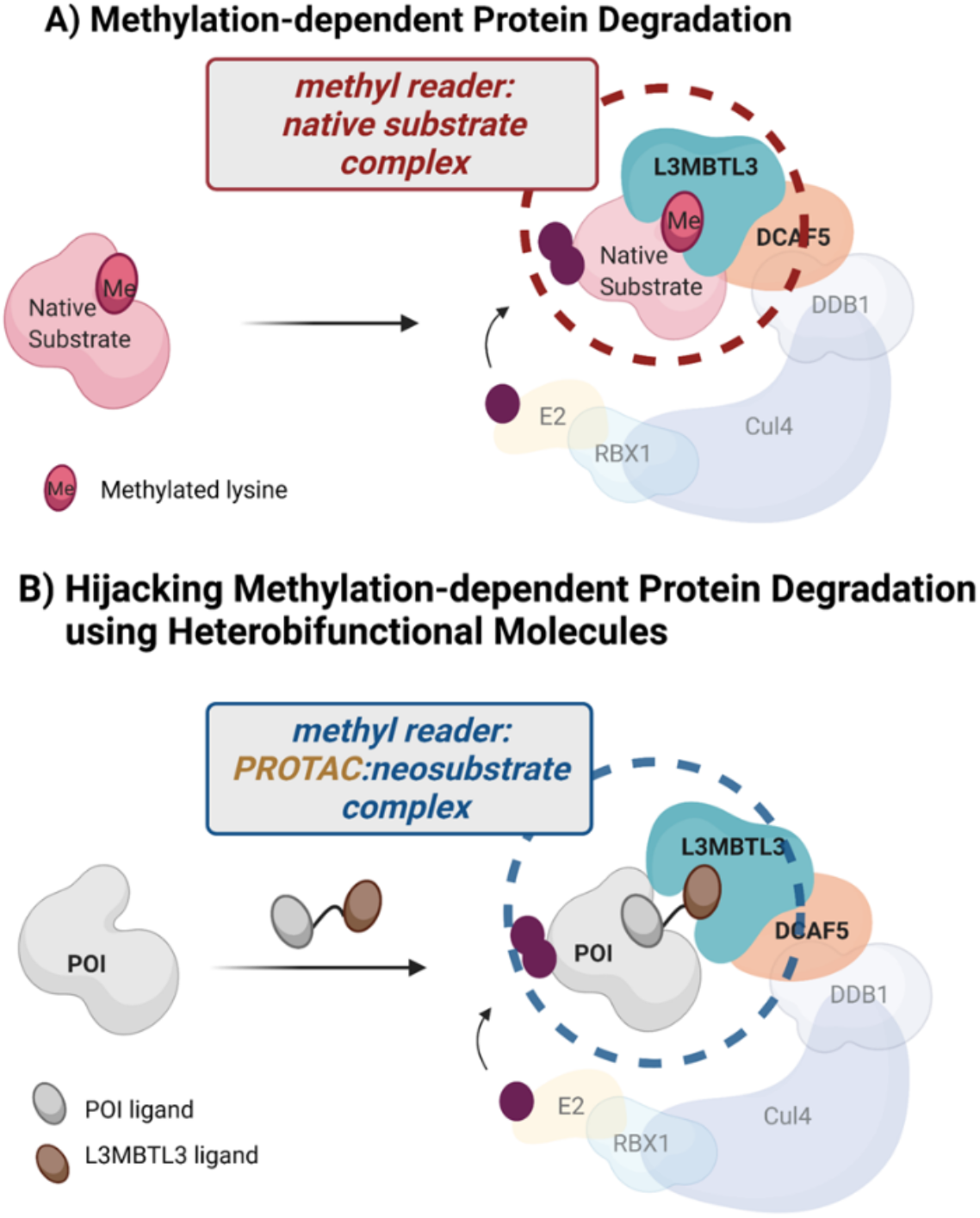
Hijacking natural methylation-dependent protein degradation using heterobifunctional molecules. A) Natural methylation-dependent protein degradation system. Methyl lysine reader protein L3MBTL3 binds to methylated proteins and targets them for proteasomal degradation via the Cul4^DCAF5^ complex. B) Hijacking methylation-dependent protein degradation using heterobifunctional molecules. L3MBTL3-recruiting PROTACs induce ternary complex formation between a protein of interest (POI) and L3MBTL3, promoting ubiquitination and degradation of the POI via the Cul4^DCAF5^ complex.

By co-opting this natural methylation-dependent protein degradation system using chimeric molecules, we demonstrate the nuclear-restricted degradation of target proteins via the recruitment of the L3MBTL3-Cul4^DCAF5^ complex. Specifically, L3MBTL3-recruiting PROTACs induce nuclear FKBP12 and BRD2 protein degradation in a time, dose, proteasome- and L3MBTL3-dependent manner. We hypothesize that this approach can be extended to other ligandable reader protein: E3 ligase complexes to expand the E3 ligase repertoire in TPD.

## Results

### L3MBTL3 ligand (UNC1215) does not disrupt L3MBTL3-Cul4^DCAF5^ interaction

Prior studies have shown that methyl reader protein L3MBTL3 complexes with Cul4^DCAF5^ to recognize and help ubiquitinate methylated proteins^30–31^. Analogous to the use of a VHL E3 ligase small molecule ligand in PROTAC design, we explored whether the methyl-lysine binding pocket of L3MBTL3 could be hijacked to induce protein of interest (POI) degradation using a heterobifunctional molecule (Figure 1, Figure S1)^6, 32^. Starting with UNC1215, a potent antagonist of L3MBTL3 that binds to its methyl-lysine binding pocket (Figure 2A), we first confirmed that UNC1215 binding to L3MBTL3 does not disrupt L3MBTL3-Cul4^DCAF5^ interaction^33^. We synthesized a previously reported biotinylated UNC1215^33^ (Figure 2B) and performed a biotin pulldown experiment in 293T cells expressing L3MBTL3-HA and myc-DCAF5. As shown in Figure 2C, biotinylated UNC1215 binding to L3MBTL3 can be competed with excess free UNC1215 ligand. Importantly, biotinylated UNC1215 did not disrupt the L3MBTL3-DCAF5 interaction, suggesting that UNC1215 is a suitable handle for functional E3 ligase recruitment in PROTAC development. We further confirmed the interaction of L3MBTL3 with the DCAF5 E3 ligase complex by immunoprecipitating L3MBTL3-HA in the absence or presence of UNC1215 in 293T and HCT116 cells (Figure 2D, 2E). Consistent with the biotin pulldown experiment, DCAF5 and DDB1 were co-immunoprecipitated with L3MBTL3 in two cell lines irrespective of the presence of UNC1215, further suggesting that it is a suitable ligand for recruiting the L3MBTL3-Cul4^DCAF5^ E3 ligase complex to a target protein.

**Figure 2.**
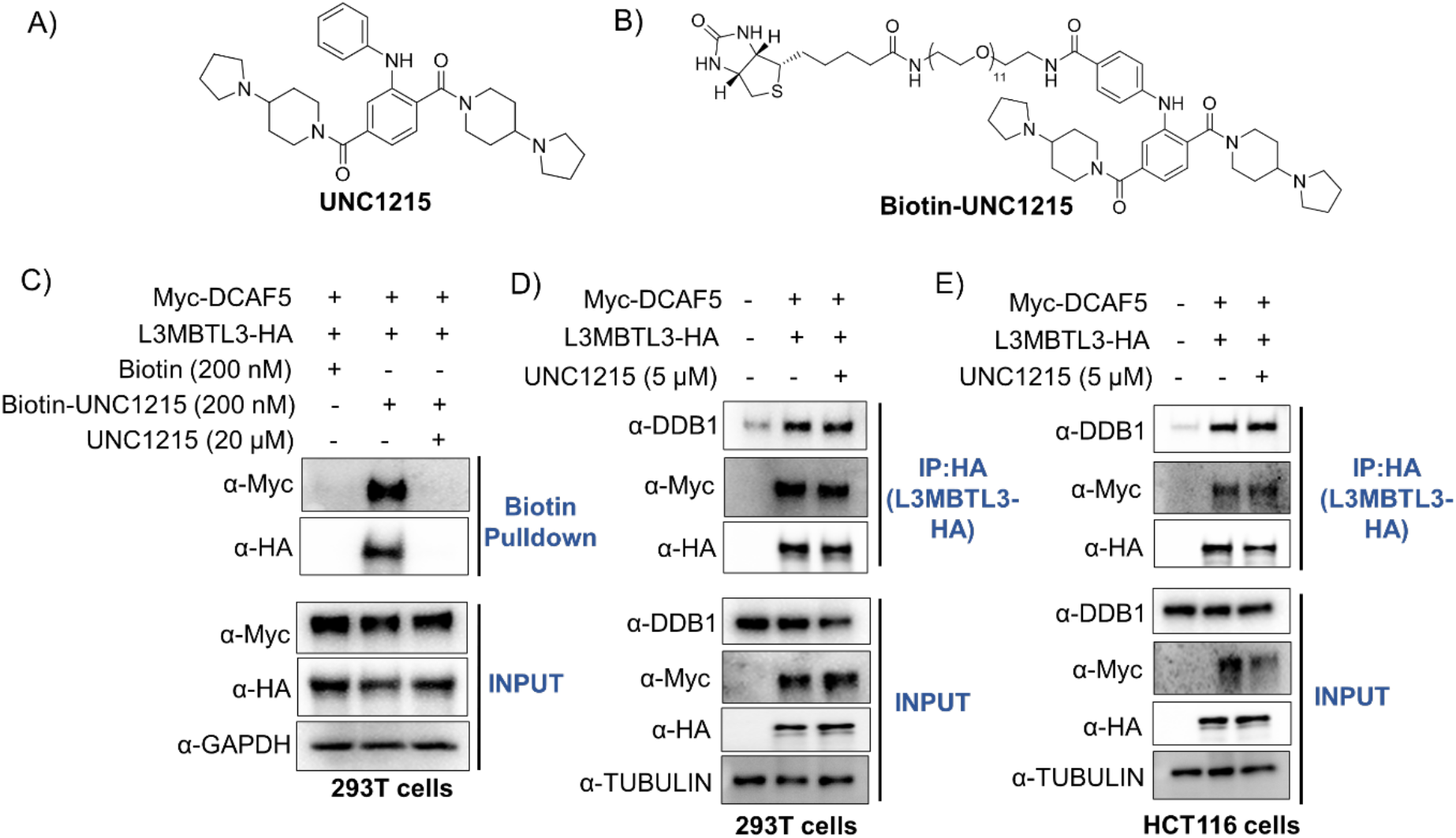
UNC1215 does not disrupt the L3MBTL3-Cul4^DCAF5^ interaction. A) Structure of the L3MBTL3 antagonist, UNC1215. B) Structure of biotin-UNC1215. C) UNC1215 does not disrupt the L3MBTL3-Cul4^DCAF5^ interaction. HEK293T (293T) cells were transiently transfected with L3MBTL3-HA and myc-DCAF5, lysed after 48 h and incubated in the presence or absence of 20 µM UNC1215 followed by biotin probe (200 nM) pulldown (n = 2). D) L3MBTL3 interacts with DCAF5 and DDB1 in HEK293T cells and E) HCT116 cells. HEK293T or HCT116 cells were co-transfected with L3MBTL3-HA and myc-DCAF5, lysed after 48 h and incubated in the presence or absence of 5 µM UNC1215 followed by immunoprecipitation of L3MBTL3-HA (n = 2).

### L3MBTL3-recruiting PROTACs induce nuclear-specific FKBP12^F36V^ degradation

Next, we wondered whether we could recruit the L3MBTL3-Cul4^DCAF5^ E3 ligase complex using heterobifunctional molecules, comprised of UNC1215 and a target protein ligand, to induce neo-substrate degradation. As a proof-of-concept, we first selected the FKBP12^F36V^ mutant protein as the target protein given the availability of a potent and selective ligand with which to evaluate this system ^34^. We attached the linker to the solvent exposed phenyl group of UNC1215, based on both the crystal structure of L3MBTL3:UNC1215 and the biotin-UNC1215 probe’s ability to bind L3MBTL3 (Figure S2). We fused a mutant-selective derivative of the synthetic ligand of FKBP12 (SLF’) to L3MBTL3-binding UNC1215 to generate SLF’-UNC1215 PROTACs with varying linker lengths (Figure 3A, Figure S3). Given the nuclear-restricted expression of L3MBTL3, we next explored whether L3MBTL3-recruiting PROTACs could induce compartment-specific target protein degradation. To this end, we generated HEK293 cells stably expressing 1) nuclear FKBP12^F36V^ [FLAG-**H**aloTag7-**F**KBP12^F36V^ with a **N**uclear **L**ocalization **S**ignal (**HFNLS**)], and 2) cytoplasmic FKBP12^F36V^ [FLAG-**H**aloTag7-**F**KBP12^F36V^ with a **N**uclear **E**xport **S**ignal (**HFNES**)] fusion proteins (Figure 3B, Figure S4). To confirm the compartment-specific expression of FKBP12^F36V^, we performed confocal microscopy analysis in each cell line via TMR chloroalkane-labeling of the HaloTag domains. As anticipated, we observed nuclear FKBP12 localization in the HFNLS cells, whereas HFNES was exclusively limited to cytoplasm, providing a system in which to assess L3MBTL3-induced, nuclear-specific protein degradation (Figure 3B). We then evaluated SLF’-UNC1215 PROTACs (KL-1 to KL-5) in both HFNLS and HFNES cell lines. KL-2 and KL-4 both induced ∼50% degradation of nuclear FKBP12 at 10 µM (Figure 3C). However, none of the SLF’-UNC1215 PROTACs could induce cytoplasmic FKBP12 degradation, indicating that L3MBTL3-recruiting PROTACs promote nuclear-specific protein degradation (Figure 3C).

**Figure 3.**
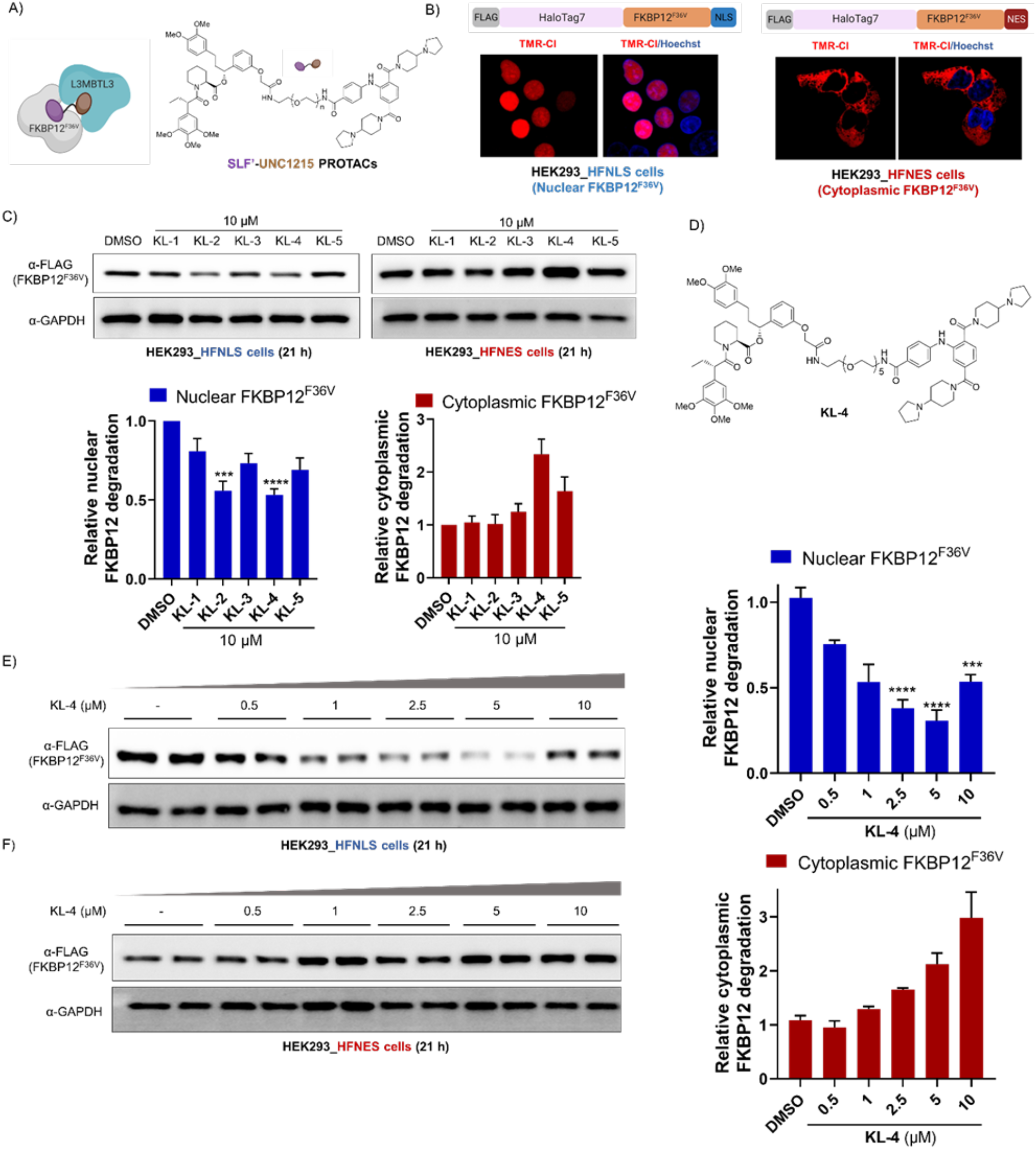
L3MBTL3-recruiting PROTACs induce nuclear-specific FKBP12^F36V^ degradation. A) Schematic of FKBP12^F36V^ and L3MBTL3 proteins in complex with SLF’-UNC1215 PROTACs. Chemical structure of SLF’-UNC1215 PROTACs (n = 2-6). B) Schematic of HFNLS and HFNES protein constructs. Confocal images of nuclear FKBP12^F36V^ and cytoplasmic FKBP12^F36V^ in HEK293 cells after labeling of HaloTag7 domain with TMR-Cl ligand. Nuclei were stained with Hoechst stain. C) SLF’-UNC1215 PROTAC screen in HEK293_HFNLS and HEK293_HFNES cell lines. HEK293 cells stably expressing nuclear FKBP12^F36V^ (HFNLS) or cytoplasmic FKBP12^F36V^ (HFNES) were treated with 10 µM of PROTACs for 21 h and subjected to western blotting. Quantitation is shown below in a bar graph format and the quantified data represent mean ± SEM (HFNLS cells, n = 4, ***P<0.001, ****P<0.0001. HFNES cells, n = 3). D) Structure of KL-4 PROTAC. E) KL-4 induces nuclear FKBP12^F36V^ degradation in a dose-dependent manner. HEK293_HFNLS cells were treated with increasing concentration of KL-4 for 21 h and blots were probed with anti-FLAG and -GAPDH antibodies. Quantitation is shown on the right and the quantified data represent mean ± SEM (n = 2, ***P<0.001, ****P<0.0001). F) KL-4 does not induce cytoplasmic FKBP12^F36V^ degradation. HEK293_HFNES cells were treated with increasing concentration of KL-4 for 21 h and then analyzed by anti-FLAG and -GAPDH antibodies. Quantitation is on the right and the quantified data represent mean ± SEM (n = 2).

Next, we selected KL-4 as the lead PROTAC to perform further experiments to validate our methyl reader-E3 ligase-based degradation system (Figure 3D). We treated both HFNLS and HFNES cell lines with increasing concentrations of KL-4, which significantly reduced nuclear FKBP12^F36V^ levels in a dose-dependent manner, with a slight hook effect detectable at 10 µM (Figure 3E). In contrast, increasing concentrations of KL-4 displayed a stabilization of cytoplasmic FKBP12^F36V^ in the HFNES cell line (Figure 3F). The stabilization of cytoplasmic FKBP12^F36V^ was also evident in free SLF’ ligand treated cells, suggesting a possible ligand-induced stabilization of FKBP12^F36V^ in the absence of induced degradation (Figure S5). Overall, the data suggest that methyl reader L3MBTL3 can be successfully recruited by the KL-4 PROTAC to induce target protein degradation in a nuclear-specific manner.

### Kinetics of KL-4 induced nuclear FKBP12 degradation

Next, we assessed the kinetics of KL-4-induced nuclear FKBP12^F36V^ degradation in HFNLS cells. HFNLS cells were treated with 10 µM of KL-4 for 3 to 24 h and evaluated for FKBP12^F36V^ degradation. KL-4 induced 50% degradation as early as 3 h, reaching 70% degradation at 6 h which is maintained up to 24 h (Figure 4A). To study the catalytic nature of L3MBTL3-recruiting PROTACs, we performed a washout experiment, where 5µM of KL-4 was incubated for 6 h and then replaced with fresh medium for 24 h, 48 h and 72 h. As the data indicate, KL-4 induced FKBP12^F36V^ degradation in both continuous 24 h treatment as well as in the 6 h washout conditions for up to 48 h (Figure 4B). As has been seen with VHL- and CRBN-recruiting PROTACs, restoration of protein levels ultimately occurs with sufficient PROTAC-free recovery time allowed ^35^. Overall, the data show that the L3MBTL3-recruiting PROTAC rapidly induces target degradation and suggests a catalytic mode of action.

**Figure 4.**
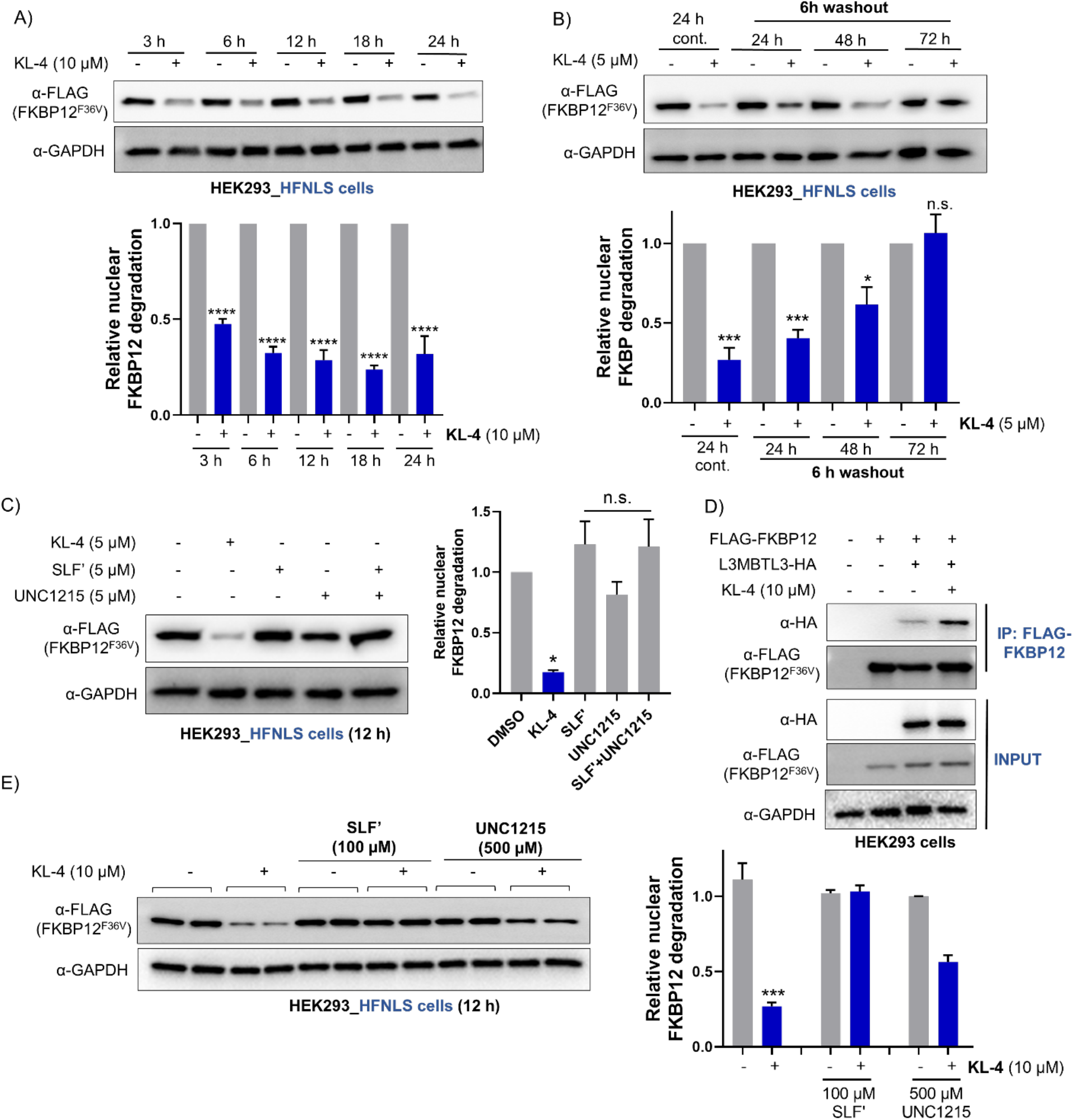
KL-4 induced nuclear FKBP12^F36V^ degradation is time-dependent and requires a ternary complex. A) Time-dependent degradation of nuclear FKBP12^F36V^ by KL-4. HEK293_HFNLS cells were treated with 10 µM KL-4 for indicated time points and analyzed for degradation by western blotting with FLAG and GAPDH antibodies. Quantitation is below and the quantified data represent mean ± SEM (n = 2, ****P<0.0001). B) L3MBTL3-recruiting PROTAC displays a catalytic mode of action. HEK293_HFNLS cells were treated with 5 µM KL-4 for 6 h followed by a washout for up to 24 h, 48 h or 72 h. Continuous 24 h treatment (24 h cont.) of 5µM KL-4 was performed as an additional control. Western blotting with FLAG antibody represents the FKBP^F36V^ protein levels in each condition. Quantitation is below and the quantified data represent mean ± SEM (n = 3, ***P<0.001, n.s: not significant). C) HEK293_HFNLS cells were treated with 5 µM KL-4, SLF’, UNC1215 or combined SLF’ and UNC1215 for 12 h and subjected to western blotting with anti-FLAG and -GAPDH antibodies. Quantitation is on the right and the quantified data represent mean ± SEM (n = 2, *P<0.05, n.s: not significant). D) FKBP12 forms a ternary complex with L3MBTL3 in a KL-4 dependent manner. HEK293_HFNLS cells were transfected with L3MBTL3-HA for 48 h and lysed. Lysates were incubated with 10 µM KL-4 for 2 h, followed by FLAG immunoprecipitation (n = 2). E) Excess SLF’ or UNC1215 abrogated KL-4 induced FKBP12^F36V^ degradation. HEK293_HFNLS cells were pre-treated with 100 µM SLF’ or 500 µM UNC1215, and then co-treated with 10 µM KL-4 for 12 h. Quantitation is on the right and the quantified data represent mean ± SEM (n = 2, ***P<0.001).

### KL-4 induced nuclear FKBP12 degradation requires a ternary complex formation

Next, the effect of the individual ligands (SLF’ or UNC1215) or combination treatment (SLF’+UNC1215) on nuclear FKBP12^F36V^ levels was assessed. As data indicate, FKBP12^F36V^ degradation was observed only with KL-4, whereas no FKBP12^F36V^ degradation was evident from treatment with ligands alone or in combination suggesting that binary complex formation is not sufficient to induce target degradation. (Figure 4C). To confirm further, we performed a ternary complex assay by transfecting L3MBTL3-HA into HFNLS cells and incubated the cell lysates in the presence or absence of 10 µM KL-4 followed by FLAG immunoprecipitation. The data indicated a significant enrichment of L3MBTL3 with FKBP12^F36V^ in the presence of KL-4 compared to DMSO-treated samples (Figure 4D). We next performed ligand competition experiments to confirm the requirement of a ternary complex for KL-4 induced FKBP12^F36V^ degradation. Pre-incubation with excess SLF’ ligand completely blocked KL-4 induced FKBP12^F36V^ degradation, whereas pre-incubation with the UNC1215 ligand partially rescued the degradation indicating that the ternary complex formation is required for the observed FKBP12^F36V^ degradation (Figure 4E).

### KL-4-mediated FKBP12 degradation is proteasome- and L3MBTL3-dependent

To assess whether FKBP12^F36V^ degradation induced by KL-4 occurs via the proteasomal pathway, HFNLS cells were treated with 10 µM KL-4 in the absence or presence of the neddylation inhibitor, MLN4924 (1 µM) or proteasome inhibitor, epoxomicin (0.5 µM). Pre-treatment with either inhibitor blocked KL-4-mediated FKBP12^F36V^ degradation, further validating its dependency on both the Cullin-RING ubiquitin ligase and the proteasome (Figure 5A). To confirm that KL-4 induced degradation occurs via L3MBTL3 protein, we took advantage of UNC1079: a previously reported negative control probe of L3MBTL3 (Figure 5B)^33^. To make the inactive control probe structurally similar to KL-4 and to keep the linker attachment position consistent, we derivatized UNC1079 by adding an aniline group: UNC1079* (Figure 5B). The only remaining structural difference between UNC1215 and UNC1079* is that the former has terminal pyrrolidine rings whereas the latter has piperidine rings. Prior to synthesizing the inactive variant of the PROTAC, we first validated the negative control probe by generating a biotinylated version, biotin-UNC1079* (Figure S6) and performed a biotin pulldown assay in 293T cells along with active biotin-UNC1215 as a positive control (Figure 2B). Interestingly, while biotin-UNC1215 consistently interacted with L3MBTL3 in a dose-dependent manner, the negative control probe, biotin-UNC1079*, did not interact with L3MBTL3 even at 5 µM (Figure 5C). Accordingly, we then generated a candidate inactive PROTAC, KL-6, by linking UNC1079* to the SLF’ ligand and evaluated the effect of KL-6 on FKBP12^F36V^ degradation (Figure 5D): KL-6 had a minimal effect on FKBP12^(F36V)^ degradation, suggesting that KL-4 induced FKBP12^F36V^ degradation is L3MBTL3-dependent (Figure 5E).

**Figure 5.**
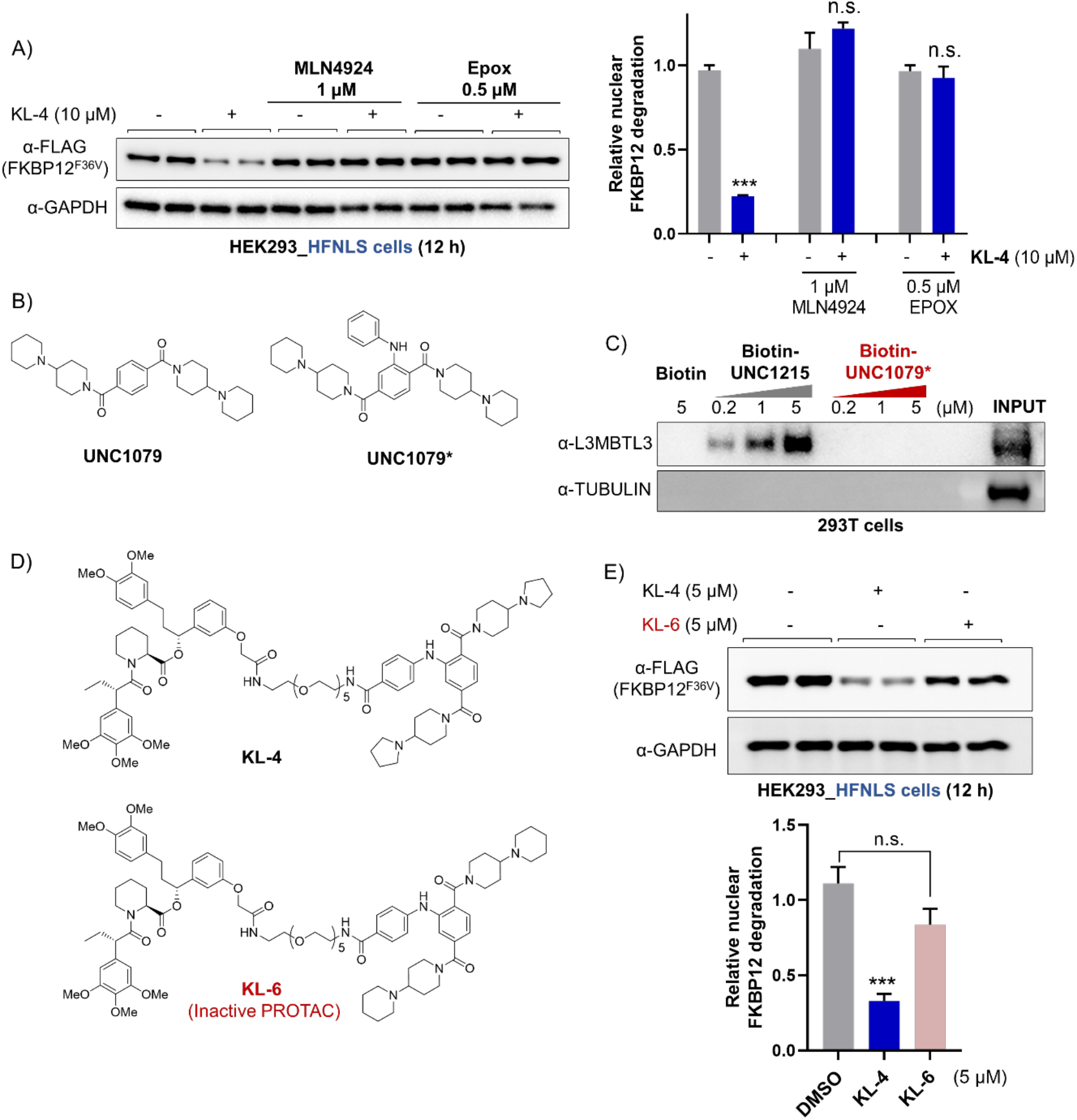
KL-4-mediated FKBP12^F36V^ degradation is proteasome- and L3MBTL3-dependent. A) KL-4 induced **FKBP12^F36V^** degradation is neddylation- and proteasome-dependent. HEK293_HFNLS cells were pre-treated with 1 µM MLN4924 or 0.5 µM epoxomicin for 1.5 h, and then co-treated with 10 µM KL-4 for 12 h. Quantitation is on the right and the quantified data represent mean ± SEM (n = 2, ***P<0.001, n.s: not significant). B) Chemical structures of UNC1079 and UNC1079*. C) Biotin pulldown assay for capture of endogenous L3MBTL3 using active (biotin-UNC1215) and inactive (biotin-UNC1079*) biotin probes. Cell lysates were incubated with increasing concentrations (0.2-5 µM) of biotin probes bound to streptavidin beads followed by western blotting with L3MBTL3 and tubulin antibodies (n = 2). D) Chemical structures of active (KL-4) and inactive (KL-6) PROTACs. E) Inactive PROTAC KL-6 does not induce significant FKBP12 degradation compared to active PROTAC, KL-4. HEK293_HFNLS cells were treated with 5 µM KL-4 and KL-6 for 12 h. Quantitation is on the right and the quantified data represent mean ± SEM (n = 2, ***P<0.001, n.s: not significant).

### L3MBTL3-recruiting PROTAC, KL-7 promotes proteasomal degradation of BRD2

To extend the applicability of L3MBTL3-mediated targeted protein degradation to an endogenous nuclear protein, we selected BRD family proteins as the second POI (Figure 6A). We coupled the widely used BRD targeting ligand, JQ1, to UNC1215 via a polyethylene glycol (PEG) linker to generate KL-7 (Figure 6A)^36^. Interestingly, KL-7 induced BRD2 degradation in multiple cell lines such as HCT116, THP1 and 293T in a concentration-dependent manner, while sparing BRD4 (Figure 6B and Figure S7-S9)^37^. We then performed a ternary complex assay by co-transfecting GFP-BRD2-V5 and L3MBTL3-HA into 293T cells and then incubating them in the presence or absence of 5µM KL-7, followed by V5 immunoprecipitation. The data indicated that L3MBTL3 interacted with BRD2 in a KL-7 PROTAC-dependent manner (Figure 6C). Next, we performed a ligand competition experiment by pre-treating HCT116 cells with excess JQ1 in the presence or absence of 5 µM KL-7. As expected, KL-7 induced BRD2 loss is blocked by excess JQ1 ligand treatment, confirming the need for a full ternary complex to achieve effective degradation (Figure 6D). To further investigate whether KL-7 induced BRD2 degradation occurs via a proteasomal pathway, HCT116 cells were pre-treated with neddylation or proteasome inhibitors in the absence or presence of 5 µM KL-7. The data indicated that KL-7 induced BRD2 degradation is prevented by both MLN4924 and epoxomicin, further reinforcing that KL-7 induced target degradation, like that of KL-4, is Cullin RING ligase- and proteasome-dependent (Figure 6E).

**Figure 6.**
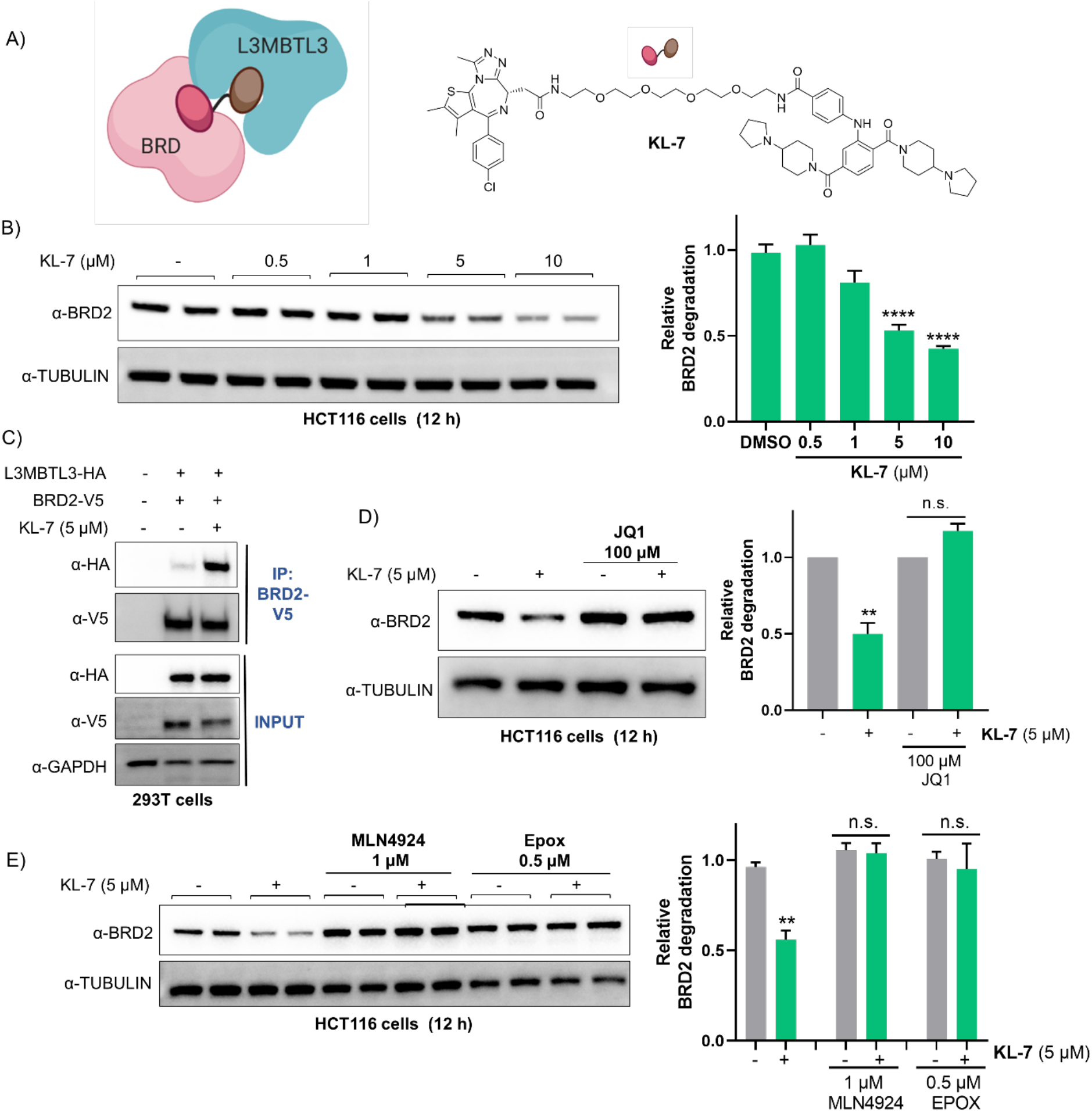
L3MBTL3-recruiting PROTAC (KL-7) promotes proteasomal degradation of BRD2. A) Schematic of BRD and L3MBTL3 proteins in complex with the JQ1-UNC1215 PROTAC. Structure of KL-7. B) KL-7 degrades BRD2 in a dose-dependent manner in HCT116 cells. HCT116 cells were treated with increasing concentrations of KL-7 for 12 h. Quantitation is on the right and the quantified data represent mean ± SEM (n = 3, ****P<0.0001). C) BRD2 forms a ternary complex with L3MBTL3 in a KL-7 dependent manner. HEK293T cells were co-transfected with L3MBTL3-HA and GFP-BRD2-V5 for 48 h and lysates were incubated with 5 µM KL-7 for 2 h, followed by V5 immunoprecipitation (n = 2). D) Excess JQ1 ligand blocks KL-7 induced BRD2 degradation. HCT116 cells were pre-treated 100 µM JQ1 for 1.5 h, and then co-treated with 5 µM KL-7 for 12 h. Quantitation is on the right and the quantified data represent mean ± SEM (n = 2, **P<0.005). E) KL-7 induced BRD2 degradation is neddylation and proteasome-dependent. HCT116 cells were pre-treated with 1 µM MLN4924 and 0.5 µM epoxomicin for 1.5 h, and then co-treated with 5 µM KL-7 for 12 h. Quantitation is on the right and the quantified data represent mean ± SEM (n = 2, **P<0.005).

## Discussion

In this study, we show the utility of a methyl-lysine reader protein ligand in PROTAC design for the recruitment of novel E3 ligases for TPD. The methyl degron was first described in 2012, when it was shown that the E3 ligase DCAF1 binds to methylated RORα via its chromo domain and subsequently targets RORα for proteasomal degradation^25^. Similarly, WWP2 is another E3 ligase that targets the methylated protein SOX2 for degradation^29^. There are only a few E3 ligases known to display such direct specificity for methylated substrates, and further studies are needed to fully understand what fraction of E3 ligases can contribute to methylation-dependent protein degradation. In addition to direct E3 ligase recruitment of methyl degrons, recent studies have shown that methylated proteins can also be recognized via methyl-lysine reader adapter proteins that target them for proteolysis by a reader bound-E3 ligase complex^31^. Consequently, we questioned whether a heterobifunctional molecule, which we have used with great success to recruit neosubstrates directly to E3 ligases for processive degradation, could also effectively mediate neosubstrate ubiquitination upon recruitment to a methyl-lysine reader adapter protein.

Here, we focused on L3MBTL3, a methyl-lysine binding protein which is known to induce its native substrate degradation in a methylation-dependent manner^31^. As a proof-of-concept study, we took advantage of a previously reported L3MBTL3 antagonist, UNC1215, as a handle to recruit L3MBTL3-bound E3 ligase complex to the vicinity of our selected target proteins: FKBP12^F36V^ and BRD2. The FKBP12^F36V^ degradation induced by our L3MBTL3-recruiting PROTAC, KL-4, was time-, concentration- and proteasome-dependent. To validate the L3MBTL3-dependent FKBP12^F36V^ degradation, we generated an inactive PROTAC based on a previously reported variant of UNC1215 that does not interact with L3MBTL3. Interestingly, the inactive PROTAC, KL-6, did not induce significant FKBP12^F36V^ degradation compared to KL-4. In the case of both our FKBP12-targeting and BRD-targeting PROTACs, recruitment of L3MBTL3 resulted in target degradation that proceeded via the characteristic mechanism of PROTACs, as assessed through use of proteasome pathway inhibitors. Furthermore, we show that the BRD-targeting, L3MBTL3-recruiting PROTAC (KL-7) display selective degradation of BRD2 over BRD4 among multiple cell lines, which was not commonly observed with VHL- and CRBN-recruiting PROTACs^37–39^. Given that no structural information of this ternary complex is available at this point, we hypothesize that this could be due to differential orientations of recruited BRD2 and BRD4 proteins with respect to the E2 bound-L3MBTL3-Cul4^DCAF5^ complex.

The L3MBTL3-recruiting PROTACs demonstrate nuclear-specific protein degradation owing to the nuclear localization of L3MBTL3. The compartment-specific degradation is a key advantage of this L3MBTL3-DCAF5 system compared to the existing, commonly utilized E3 ligases such as VHL and CRBN, which induce degradation of both the cytoplasmic and nuclear pools of target proteins. Higher potency and selectivity can be achieved by developing a PROTAC that binds to both the E3 ligase and the target protein within the same subcellular compartment. Among recently harnessed E3 ligases for TPD, only DCAF16 showed compartment-specific degradation^16^. However, the DCAF16-recruiting ligand, KB02 is a highly reactive electrophilic scout fragment which can modify multiple proteins inside the cell, limiting its broader utility in PROTAC development. Alternatively, the L3MBTL3-recruiting reversible ligand, UNC1215, can be utilized to develop PROTACs that target nuclear oncogenic proteins such as transcription factors to achieve nuclear-specific degradation with less risk of broader off-target effects that use of KB02 might incur due to the latter’s binding partner promiscuity.

In summary, we have co-opted methylation-dependent protein degradation using chimeric molecules and demonstrated the utility of a methyl reader-bound E3 ligase complex in TPD. To further understand the generalizability of this approach, we are currently exploring other reader: E3 ligase complexes that are amenable to ligand-induced protein degradation. Discovering ligands for E3 ligases with tumor-specific, tissue-specific and compartment-specific properties would address limitations such as on-target toxicity and emerging resistance to the widely used VHL- and CRBN-recruiting PROTACs^14^. We propose that ligandable tissue- or tumor-specific methyl reader proteins can be exploited in PROTAC design as an alternative to the direct recruitment of E3 ligases. While we demonstrate the proof-of-concept study by showing the utility of methyl reader proteins as E3 recruiting handles for the first time in TPD, we anticipate that this will be a new direction to expand the E3 ligase toolbox to achieve compartment- and disease-specific protein degradation in the future.

## Acknowledgements

We would like to thank Dr. Kusal Samarasinghe, Dr. Saul J. Figueroa, Dr. Mackenzie Krone and Michael Bond for their helpful suggestions and critiques. We would like to thank Dr. Kanak Raina for providing HFNLS and HFNES cell lines. C.M.C. is funded by the NIH (R35CA197589 and R01CA238570) and is supported by an American Cancer Research Professorship. Figures were created with BioRender.com

## Author contributions

D.A.N. and C.M.C. conceived the research project. **^#^**D.A.N and K.L. contributed equally to this work. D.A.N. designed and performed biological experiments and wrote the manuscript with input from all the co-authors. K.L. designed, synthesized and characterized the compounds.

## Methods-Biology

### Common reagents and antibodies

Antibodies against HA (3724), FLAG (14793), myc (2276), BRD4 (13440), V5 (13202) and GAPDH (2118) were purchased from Cell Signaling Technology. DDB1 antibody (ab109027) was purchased from Abcam. The BRD2 (A700-008) and L3MBTL3 (A302-852A) antibodies were purchased from Bethyl labs. Alexa Fluor 488 conjugated tubulin antibody (16-232) was purchased from Millipore. Alexa Fluor 488-conjugated goat anti-rabbit secondary antibody (A11008) and Hoechst 33342 stain (H3570) were purchased from Invitrogen. Amersham ECL Rabbit IgG, HRP-linked whole secondary Ab (from donkey) (NA934) and Amersham ECL Mouse IgG, HRP-linked whole secondary Ab (from sheep) (NA931) were purchased from Cytiva. Pierce™ Anti-HA Magnetic Beads (88836) and Pierce™ Streptavidin Magnetic Beads (88816) and RIPA Lysis and Extraction Buffer (89900) were purchased from Thermo Scientific. Anti-V5 magnetic beads (M167-11) obtained from MBL international corporation. Anti-FLAG M2 magnetic beads (M8823) were purchased from Sigma Aldrich. ECL™ Prime (GERPN2232) and Super Signal West Femto reagents (34095) were purchased from Millipore Sigma and Thermo Scientific™ respectively. Halotag® TMR ligand (G8251) was obtained from Promega. Poly-L-lysine solution (P8920) was purchased from Sigma Aldrich. μ-slide 8 well chambered coverslips (80826) were obtained from Ibidi. MLN4924 (S7109) was purchased from Selleck Chemicals. L3MBTL3-HA (HG15231-CY) and untagged DCAF5 (HG22894-UT) plasmids were obtained from Sinobiologicals and GFP-BRD2-V5 plasmid (Addgene #65376) was obtained from Addgene. FLAG-HFNLS and FLAG-HFNES genes containing pCDH-CMV-MCS-EF1a-plasmids were synthesized at Epoch life science.

### Cell lines

HEK293T cells, THP1 and HCT116 cells were purchased from ATCC. HEK293T cells and HCT116 cells were cultured in DMEM with 10% FBS, Penicillin (100 U/mL) and Streptomycin (100 µg/mL). HEK293 cells stably expressing HFNLS and HFNES were cultured in DMEM with10% FBS, Penicillin (100 U/mL), Streptomycin (100 µg/mL) and Puromycin (1 **μ**g/ml). THP1 cells were cultured in RPMI-1640 medium supplemented with 10% FBS, Penicillin (100 U/mL), Streptomycin (100 µg/mL), and 0.02% BME. All cells were grown in a humidified atmosphere at 37 °C and 5% CO_2_.

### Cloning

To generate the tagged version of DCAF5, myc tag was cloned into the N-terminus of DCAF5 gene via USER cloning. Human DCAF5 cDNA was PCR amplified from pCMV vector containing untagged DCAF5 (Sinobiologicals) and subcloned into the pcDNA5 FRT vector. Final constructs were sequence verified by Sanger sequencing. Full sequence information for FLAG-halotag-FKBP12^F36V-^NLS (HFNLS) and FLAG-halotag-FKBP12^F36V-^NES (HFNES) constructs is provided below.

### Generation of HFNLS and HFNES cell lines

FLAG-halotag-FKBP12^F36V-^NLS (HFNLS) and FLAG-halotag-FKBP12^F36V-^NES (HFNES) lentivirus was generated by lentiviral transduction into HEK293 cells. Then, virus containing media was collected after 48 h, filtered, and HEK293 cells were infected with collected virus in the presence of polybrene (10 µg/mL). Next, puromycin (1 µg/mL) was added after 48 h and maintained until colonies were performed. Single colonies were picked and grown separately and later validated by western blotting to confirm the expression of FLAG-halotag-FKBP12^F36V^ and by fluorescence imaging.

### Cell lysis and Western Blotting

After desired treatments, cells were collected and lysed in RIPA buffer which was supplemented with 1X EDTA-free complete protease inhibitor cocktail (Roche). Protein concentration was determined by Pierce™ BCA Protein Assay kit (23225). Proteins were separated by SDS-PAGE, transferred to PVDF membrane, and blocked with 5% (w/v) non-fat dry milk in TBST (0.05%Tween-20, 20mM Tris-HCl pH 7.6, 150mM NaCl) for 1 h at RT. The membrane was incubated with primary antibodies against FLAG, GAPDH, BRD2, BRD4, HA, myc, DDB1 and L3MBTL3 (1:10000 dilution for DDB1 and 1:1000 dilution for others) overnight at 4 °C. Following three washes with TBST, secondary anti-rabbit HRP antibody (1:5000 dilution in 5% milk in TBST) was incubated for 1 h at RT. After washing with TBST, membrane was developed using ECL or Super Signal West Femto substrate (ThermoFisher). For FLAG and GAPDH blots, Alexa Fluor 488-conjugated goat anti-rabbit secondary antibody (1:1000 dilution in 5% milk in TBST) was used. Images were recorded using ChemiDoc MP Imaging system (BioRad).

### Biotin pulldown assays

For biotin pulldown assay with overexpressed L3MBTL3 and DCAF5, L3MBTL3-HA and myc-DCAF5 plasmids were co-transfected into HEK293T cells using PEI transfection reagent. After 48 h, cells were collected, lysed (NP-40 lysis buffer: 25 mM Tris-HCl, pH 7.5, 150 mM NaCl, 5 mM MgCl2, 1% NP40, 5% glycerol+ 1X protease inhibitor) and protein concentration was determined by BCA assay. First, 1 mg of lysates were pre incubated with excess UNC1215 (20 µM) for 2 h at 4 °C. Biotin-probe (200 nM) was added to Streptavidin magnetic beads (30 µL) in TBST and Incubated at RT for 1 h. Following, incubation, excess biotin was removed by washing the beads twice with 1 mL of TBS. Then preincubated lysates were added to biotin-probe conjugated beads and incubated overnight at 4 °C. Next day, flowthrough was collected, and beads were washed 3 times with 500 µL of IP wash buffer (25 mM Tris-HCl pH 7.5, 150 mM NaCl, 0.4% NP40). Bound proteins were eluted with 2X sample gel loading buffer (diluted from 4X NuPAGE™ LDS sample buffer) via boiling for 8 minutes and separated by SDS-PAGE, followed by Western blot analysis. For endogenous L3MBTL3 pulldown using biotin-UNC1215 and biotin-UNC1079* probes, HEK293T cells were collected and lysed in NP40 lysis buffer. Increasing concentrations of biotin probes (0.2, 1, 5 µM) were incubated with Streptavidin magnetic beads (30 µL) in TBST and Incubated at RT for 1 h. Following incubation, excess biotin was removed by washing beads twice with 1 mL of TBS. Then lysates were added to biotin-probe modified beads and incubated overnight at 4 °C. Next day, beads were washed as described above and performed SDS-PAGE and Western blotting with appropriate antibodies.

### Co-Immunoprecipitations

For co-immunoprecipitation assay with overexpressed L3MBTL3 and DCAF5, L3MBTL3-HA and myc-DCAF5 plasmids were co-transfected into either HEK293T or HCT116 cells using PEI transfection reagent. After 48 h, cells were harvested and lysed in NP-40 lysis buffer and protein concentrations were determined by BCA assay. For the Co-Immunoprecipitation experiments, cell lysates were first incubated with DMSO or UNC1215 for 2h at 4 °C. Then lysates were incubated with pre-washed anti-HA magnetic beads overnight at 4 °C. Following day, HA-affinity beads were washed 3 times with immunoprecipitation (IP) wash buffer (25 mM Tris-HCl pH 7.5, 150 mM NaCl, 0.4% NP40) and bound proteins were eluted by boiling in 2X sample gel loading buffer. Samples were separated by SDS-PAGE, followed by Western blot analysis.

### Ternary complex assays

For the ternary complex assays with FKBP12 and L3MBTL3, HFNLS cells were transfected with L3MBTL3-HA plasmids using PEI transfection reagent. After 48 h, cells were collected and lysed in NP40 lysis buffer and lysates were first pre-incubated with DMSO or 10 µM KL-4 for 2 h at 4 °C. Then, lysates were added to anti-FLAG magnetic beads and incubated overnight at 4 °C. Next day, the beads were washed three times with mild IP wash buffer (25 mM Tris-HCl pH 7.5, 150 mM NaCl, 0.2% NP40) and bound proteins were eluted by heating at 95 °C for 10 min. The samples were then subjected to SDS-PAGE and Western blotting. For the ternary complex assays with BRD2 and L3MBTL3, HEK293T cells were co-transfected with L3MBTL3-HA and GFP-BRD2-V5 plasmids using PEI transfection reagent. Following cell lysis, lysates were pre-incubated with DMSO or 5 µM KL-7 for 2 h at 4 °C, followed by incubating with anti-V5 magnetic beads overnight at 4 °C. Next day, anti-V5 magnetic beads were washed three times with mild IP buffer (25 mM Tris-HCl pH 7.5, 150 mM NaCl, 0.2% NP40) and bound proteins were eluted using 2X sample gel loading buffer and subjected to Western blot analysis.

### Confocal microscopy analysis of HFNLS and HFNES cell lines

To validate the nuclear and cytoplasmic expression of FKBP12, HFNLS and HFNES cells were analyzed by confocal microscopy. HFNLS and HFNES cells were seeded on 8 well chambered poly-D-lysine coated coverslips (Ibidi). Next day, to visualize the HaloTag protein, cells were incubated with 500 nM HaloTag® TMR ligand (Promega) for 15 min at 37 °C. Then, cells were washed with serum containing media for 30 min at 37 °C, followed by three PBS washes. Cells were fixed with 4% paraformaldehyde in PBS for 15 min at RT. Then, cells were washed 2X with PBS and stained with Hoechst stain (3 µg/mL) for 10 min at RT followed by two PBS washes. Then, the cells were imaged using Zeiss LSM 880 confocal microscope.

### Statistical analysis

For all biological assays, at least two independent experiments were performed. The number of independent replicates, quantitation, error bars and p-values are shown in each figure legend. Images were quantified using Image Lab software (Biorad) and graphs were plotted using GraphPad Prism 9.2.0. One-way ANOVAs or two-way ANOVAs were utilized for comparison between groups. P<0.05 is considered statistically significant.

## Supporting Information

### Supplementary Note 1. Synthetic Procedures

#### Synthesis of UNC1215

**Scheme 1.**
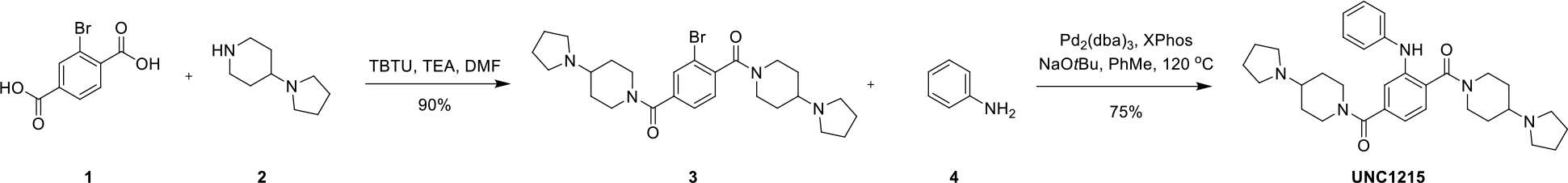
Synthesis of UNC1215

**[3-bromo-4-(4-pyrrolidin-1-ylpiperidine-1-carbonyl)phenyl]-(4-pyrrolidin-1-yl-1-piperidyl)methanone (3)**

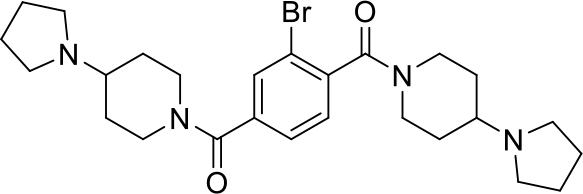

To a mixture of 2-bromoterephthalic acid (**1**) (200 mg, 0.816 mmol) and TBTU (681 mg, 2.12 mmol) in DMF (2 mL), a solution of 4-pyrrolidin-1-ylpiperidine (**2**) (290 mg, 1.88 mmol) and TEA (0.683 mL, 4.90 mmol) in DMF (1 mL) was added. The mixture was stirred at rt for 12h. The reaction was quenched by the addition of saturated aq. NaHCO_3_ (10 mL) and extracted with CH_2_Cl_2_ (15 mL x 3). The combined organic extracts were washed with sat. aq. NaHCO_3_, dried over Na_2_SO_4_ and concentrated under reduced pressure. The residue was purified by Prep-TLC (Magic base/CH_2_Cl_2_ = 3:1, magic base = CH_2_Cl_2_/MeOH/NH_3_ H_2_O = 60/10/1) to afford [3-bromo-4-(4-pyrrolidin-1-ylpiperidine-1-carbonyl)phenyl]-(4-pyrrolidin-1-yl-1-piperidyl)methanone (**3**) (382 mg, 0.738 mmol, yield: 90.4 %). ^1^H NMR (600 MHz, CD_3_OD) δ 7.73 (d, J = 11.8 Hz, 1H), 7.54 – 7.35 (m, 2H), 4.71 – 4.55 (m, 2H), 3.70 (d, J = 13.8 Hz, 1H), 3.44 (t, J = 14.6 Hz, 1H), 3.23 – 3.08 (m, 2H), 2.97 – 2.85 (m, 2H), 2.74 – 2.64 (m, 8H), 2.48 – 2.40 (m, 2H), 2.16 – 2.07 (m, 2H), 2.00 – 1.90 (m, 2H), 1.90 – 1.80 (m, 8H), 1.74 – 1.35 (m, 4H). ^13^C NMR (151 MHz, CD_3_OD, rotamers observed) δ 169.6, 168.8, 168.6, 140.6, 140.4, 139.7, 139.6, 132.3, 132.2, 129.2, 129.1, 127.5, 127.5, 120.3, 63.1, 63.0, 63.0, 52.5, 52.5, 47.7, 47.2, 46.6, 42.2, 41.6, 41.5, 32.6, 32.5, 32.2, 31.7, 31.6, 24.0, 24.0. HRMS (ESI) [M+H]^+^ for C_26_H_38_BrN_4_O_2_: 519.2158, found: 519.2156.

**UNC1215**

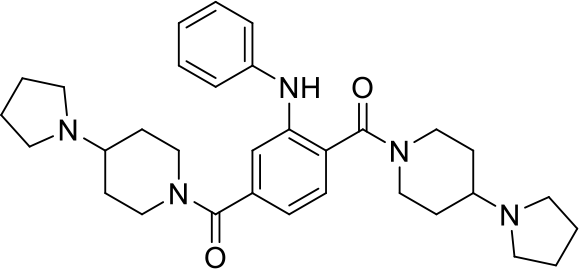

A mixture of [3-bromo-4-(4-pyrrolidin-1-ylpiperidine-1-carbonyl)phenyl]-(4-pyrrolidin-1-yl-1-piperidyl)-methanone (**3**) (25.0 mg, 0.0483 mmol), aniline (**4**) (9.00 mg, 0.0966 mmol), Pd_2_(dba)_3_ (8.85 mg, 0.00966 mmol), 2-Dicyclohexylphosphino-2’,4’,6’-tri-iso-propyl-1,1’-biphenyl (9.21 mg, 0.0193 mmol) and Sodium tert-butoxide (6.96 mg, 0.0725 mmol) in anhydrous toluene (0.5 mL) was added to a sealable reaction tube. The solution was degassed for 20 minutes and stirred at 120 °C for 15h. The reaction was cooled to room temperature, diluted with CH_2_Cl_2_, and filtered over a pad of celite. The solvent was removed under reduced pressure. The residue was purified by prep-TLC (Magic base/CH_2_Cl_2_ = 3:1, magic base = CH_2_Cl_2_/MeOH/NH_3_ H_2_O = 60/10/1) to give desired compound **UNC1215** (19.3 mg, 0.0364 mmol, yield: 75.4 %). ^1^H NMR (600 MHz, CD_3_OD) δ 7.32 (d, J = 7.8 Hz, 1H), 7.27 (t, J = 7.7 Hz, 2H), 7.21 (s, 1H), 7.10 (d, J = 7.9 Hz, 2H), 7.00 – 6.94 (m, 2H), 4.59 (d, J = 13.3 Hz, 2H), 3.81 (d, J = 13.9 Hz, 2H), 3.12 (t, J = 13.5 Hz, 2H), 2.85 (t, J = 13.2 Hz, 2H), 2.80 – 2.64 (m, 8H), 2.57 – 2.43 (m, 2H), 2.16 – 1.76 (m, 12H), 1.57 – 1.34 (m, 4H). ^13^C NMR (151 MHz, CD_3_OD) δ 171.4, 170.1, 143.6, 143.3, 139.0, 130.5, 130.0, 127.3, 123.3, 120.6, 119.3, 116.5, 63.0, 63.0, 52.5, 52.4, 47.6, 42.0, 32.5, 31.5, 24.0, 24.0. HRMS (ESI) [M+H]^+^ for C_32_H_44_N_5_O_2_: 530.3495, found: 530.3489.

#### Synthesis of Biotin-UNC1215

**Scheme 2.**
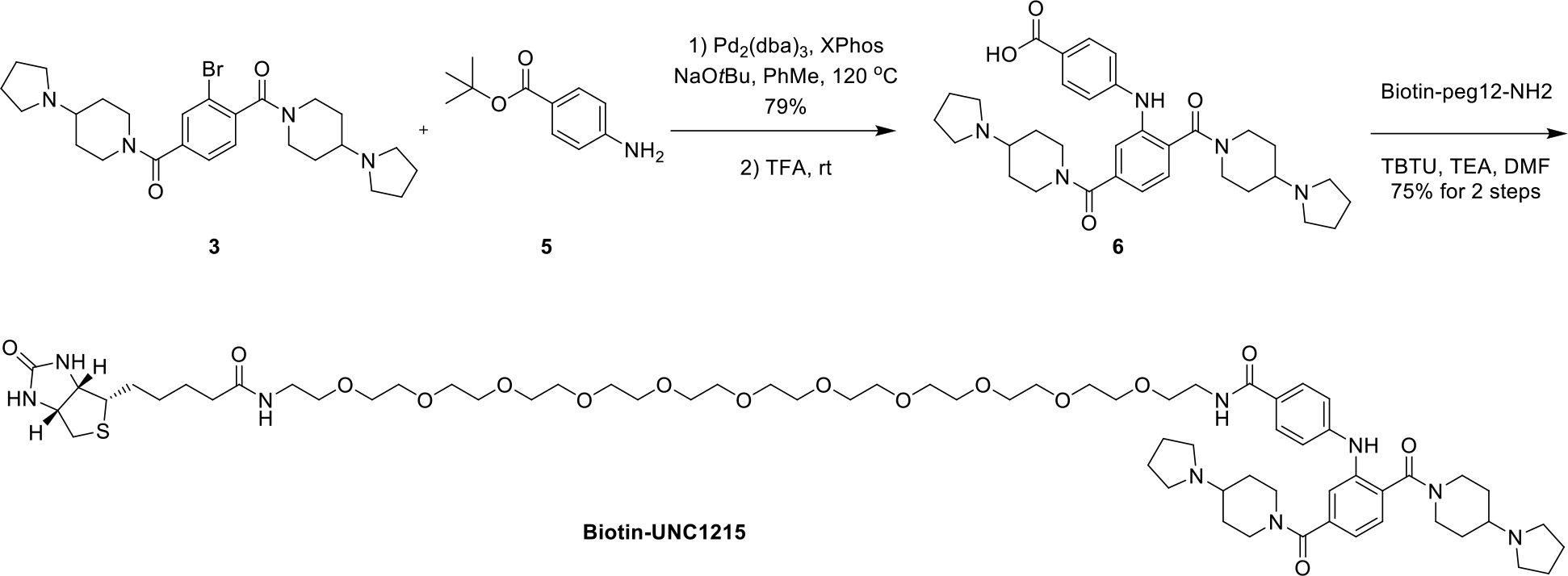
Synthesis of Biotin-UNC1215

**4-[2,5-bis(4-pyrrolidin-1-ylpiperidine-1-carbonyl)anilino]benzoic acid (6)**

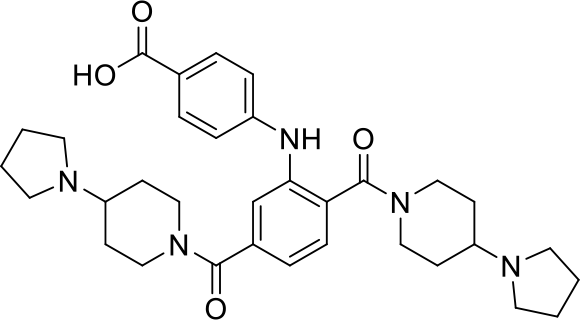

A mixture of [3-bromo-4-(4-pyrrolidin-1-ylpiperidine-1-carbonyl)phenyl]-(4-pyrrolidin-1-yl-1-piperidyl)-methanone (**3**) (104 mg, 0.201 mmol), 2-Dicyclohexylphosphino-2’,4’,6’-tri-iso-propyl-1,1’-biphenyl (38.3 mg, 0.0804 mmol), Pd_2_(dba)_3_ (36.8 mg, 0.0402 mmol), and sodium tert-butoxide (29.0 mg, 0.301 mmol) in anhydrous toluene (1 mL) was added to a sealable reaction tube. The solution was degassed for 20 minutes and tert-butyl 4-aminobenzoate (**5**) (58.3 mg, 0.301 mmol) was added subsequently. The reaction tube was tightly sealed and the reaction was stirred at 120 °C for 15h. The reaction was cooled to room temperature, diluted with CH_2_Cl_2_, and filtered over a pad of celite. The CH_2_Cl_2_ was removed under reduced pressure. The residue was purified by prep-TLC (Magic base/CH_2_Cl_2_ = 3:1, magic base = CH_2_Cl_2_/MeOH/NH_3_ H_2_O = 60/10/1) to give desired compound tert-butyl 4-[2,5-bis(4-pyrrolidin-1-ylpiperidine-1-carbonyl)anilino]benzoate (100 mg, 0.159 mmol, yield: 79.0 %). ^1^H NMR (600 MHz, CD_3_OD) δ 7.84 – 7.79 (m, 2H), 7.41 (d, J = 7.8 Hz, 1H), 7.38 (d, J = 1.5 Hz, 1H), 7.15 (dd, J = 7.8, 1.5 Hz, 1H), 7.04 (d, J = 8.6 Hz, 2H), 4.61 (d, J = 15.0 Hz, 2H), 3.87 – 3.63 (m, 2H), 3.20 – 3.00 (m, 2H), 2.93 – 2.57 (m, 10H), 2.53 – 2.39 (m, 2H), 2.16 – 1.71 (m, 12H), 1.57 (s, 9H), 1.52 – 1.26 (m, 4H). ^13^C NMR (151 MHz, CD_3_OD) δ 171.0, 169.6, 167.3, 149.2, 140.6, 139.2, 132.2, 130.2, 124.6, 122.2, 120.5, 116.8, 81.6, 81.6, 63.1, 62.8, 52.5, 52.2, 47.6, 47.3, 42.1, 41.7, 32.7, 32.1, 31.7, 31.3, 28.5, 24.0, 24.0. HRMS (ESI) [M+H]^+^ for C_37_H_52_N_5_O_4_: 630.4019, found: 630.4014.

5 mg of the obtained product was dissolved in 1 mL of TFA/CH_2_Cl_2_ (v/v = 1:2). The mixture was stirred at rt for 2h. Concentrated and the crude product (**6**) was used into next step without further purification.

**Biotin-UNC1215**

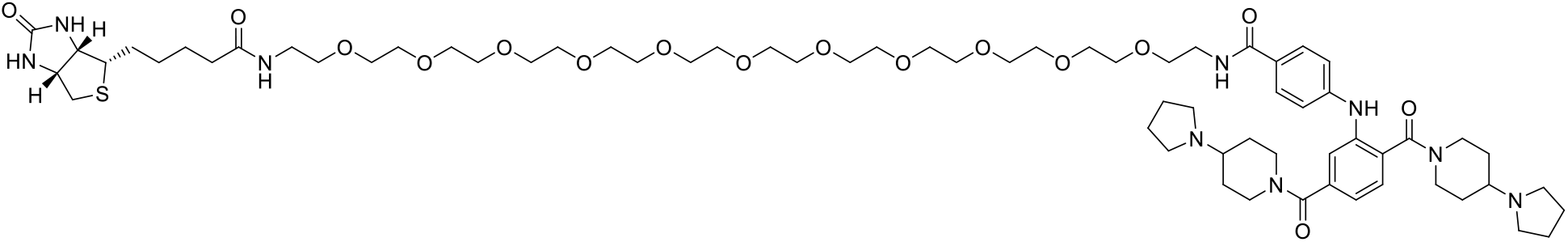

To a mixture of 4-[2,5-bis(4-pyrrolidin-1-ylpiperidine-1-carbonyl)anilino]benzoic acid (**6**) (4.50 mg, 0.00784 mmol) and TBTU (3.02 mg, 0.00941 mmol) in DMF (0.3 mL), a solution of Biotin-peg12-NH_2_ (6.05 mg, 0.00784 mmol) and TEA (0.0109 mL, 0.0784 mmol) in DMF (0.2 mL) was added. The mixture was stirred at rt for 12h. The reaction was quenched by the addition of saturated aq. NaHCO_3_ (2 mL) and extracted with CH_2_Cl_2_ (3 mL x 3). The combined organic extracts were dried over Na_2_SO_4_ and concentrated under reduced pressure. The residue was purified by Prep-HPLC (0-70 MeCN/H_2_O with 0.1% FA) to give the desired compound **Biotin-UNC1215** (7.80 mg, 0.00588 mmol, yield: 75.0 %). ^1^H NMR (500 MHz, CD_3_OD) δ 8.54 (s, 2H), 7.76 (d, J = 8.7 Hz, 2H), 7.43 (d, J = 7.8 Hz, 1H), 7.38 (d, J = 1.5 Hz, 1H), 7.15 (dd, J = 7.7, 1.5 Hz, 1H), 7.09 (d, J = 8.7 Hz, 2H), 4.69 (s, 2H), 4.49 (dd, J = 7.8, 4.9 Hz, 1H), 4.30 (dd, J = 7.9, 4.5 Hz, 1H), 3.96 – 3.46 (m, 48H), 3.36 (t, J = 5.5 Hz, 2H), 3.25 – 3.02 (m, 12H), 2.95 – 2.85 (m, 2H), 2.70 (d, J = 12.7 Hz, 1H), 2.27 – 1.91 (m, 14H), 1.78 – 1.38 (m, 10H). ^13^C NMR (126 MHz, CD_3_OD) δ 176.1, 171.2, 169.8, 169.6, 166.1, 148.0, 141.3, 139.1, 130.4, 130.1, 129.9, 127.4, 121.7, 120.0, 117.5, 71.5, 71.5, 71.5, 71.5, 71.43, 71.4, 71.4, 71.3, 71.3, 70.7, 70.6, 63.4, 63.0, 62.9, 61.6, 57.0, 52.7, 52.5, 47.2, 41.7, 41.1, 40.9, 40.4, 36.7, 31.0, 29.8, 29.5, 26.9, 24.0. HRMS (ESI) [M+H]^+^ for C_67_H_108_N_9_O_16_S: 1326.7635, found: 1326.7629.

#### Synthesis of Biotin-UNC1079*

**Scheme 3.**
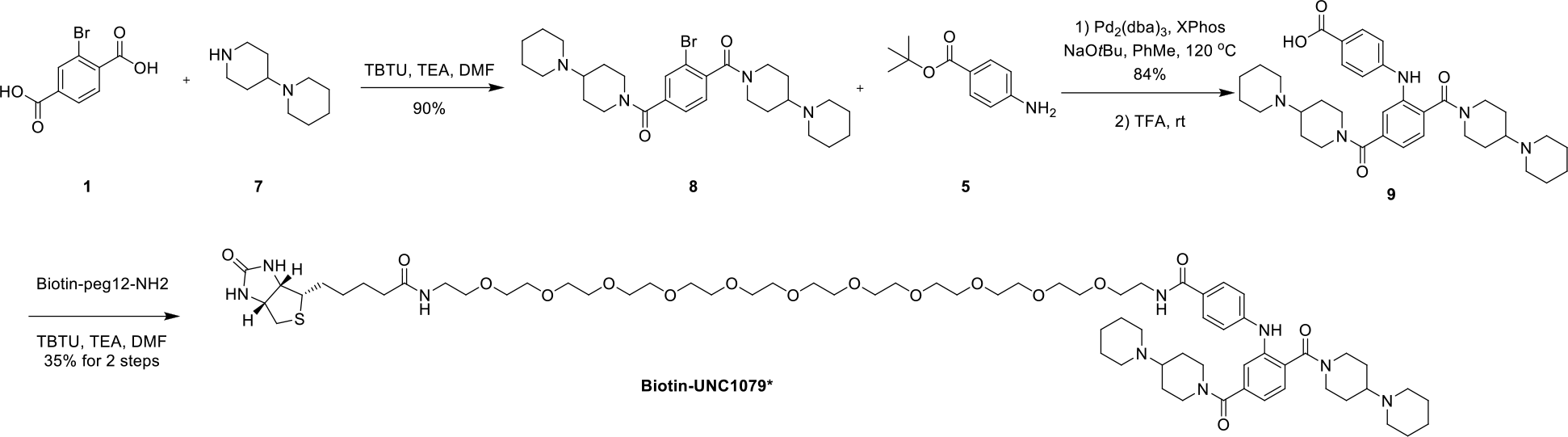
Synthesis of Biotin-UNC1079*

**[3-bromo-4-[4-(1-piperidyl)piperidine-1-carbonyl]phenyl]-[4-(1-piperidyl)-1-piperidyl]methanone (8)**

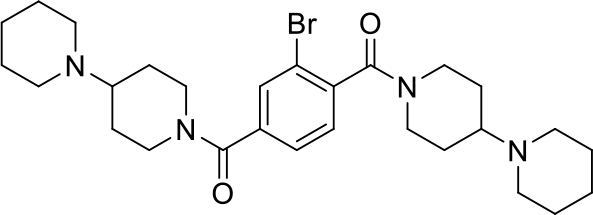

To a mixture of 2-bromoterephthalic acid (**1**) (50.0 mg, 0.204 mmol) and TBTU (170 mg, 0.531 mmol) in DMF (0.5 mL), a solution of 1-(4-piperidyl)piperidine (**7**) (79.0 mg, 0.469 mmol) and TEA (0.171 mL, 1.22 mmol) in DMF (0.3 mL) was added. The mixture was stirred at rt for 12h. The reaction was quenched by the addition of saturated aq. NaHCO_3_ (10 mL) and extracted with CH_2_Cl_2_ (15 mLx3). The combined organic extracts were washed with sat. aq. NaHCO_3_, dried over Na_2_SO_4_ and concentrated under reduced pressure. The residue was purified by Prep-TLC (Magic base/CH_2_Cl_2_ = 3:1, magic base = CH_2_Cl_2_/MeOH/NH_3_.H_2_O = 60/10/1) to afford [3-bromo-4-[4-(1-piperidyl)piperidine-1-carbonyl]phenyl]-[4-(1-piperidyl)-1-piperidyl]methanone (**8**) (100 mg, 0.183 mmol, yield: 89.8 %). ^1^H NMR (500 MHz, CD_3_OD) δ 7.72 (d, J = 12.2 Hz, 1H), 7.52 – 7.34 (m, 2H), 4.79 – 4.62 (m, 2H), 3.74 (d, J = 13.6 Hz, 1H), 3.52 – 3.40 (m, 1H), 3.24 – 3.05 (m, 2H), 2.94 – 2.78 (m, 2H), 2.70 – 2.50 (m, 10H), 2.08 – 1.80 (m, 4H), 1.76 – 1.38 (m, 16H). ^13^C NMR (126 MHz, CD_3_OD, rotamers observed) δ 169.6, 168.8, 168.6, 140.6, 140.4, 139.7, 139.6, 132.4, 132.2, 129.2, 129.1, 127.6, 127.5, 120.3, 120.2, 63.5, 63.5, 63.4, 51.2, 48.1, 47.4, 43.0, 42.5, 42.3, 29.4, 29.3, 29.3, 29.0, 28.6, 28.5, 28.5, 28.4, 26.8, 26.8, 25.5. HRMS (ESI) [M+H]^+^ for C_28_H_42_BrN_4_O_2_: 547.2471, found: 547.2470.

**4-[2,5-bis[4-(1-piperidyl)piperidine-1-carbonyl]anilino]benzoic acid (9)**

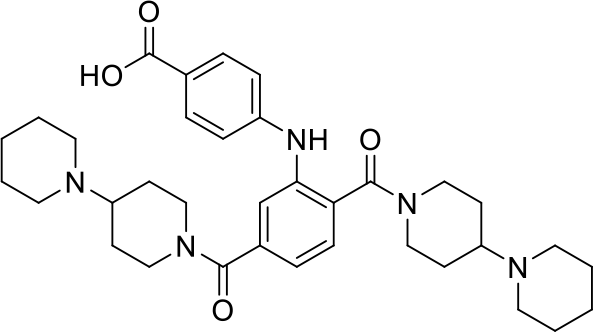

A mixture of [3-bromo-4-[4-(1-piperidyl)piperidine-1-carbonyl]phenyl]-[4-(1-piperidyl)-1-piperidyl]meth-anone (**8**) (55.0 mg, 0.101 mmol), 2-dicyclohexylphosphino-2′,4′,6′-triisopropylbiphenyl (XPhos) (24.4 mg, 50 mol%), tris(dibenzylideneacetone)dipalladium(0) (23.4 mg, 25 mol%), and Sodium tert-butoxide (14.8 mg, 0.154 mmol) in anhydrous toluene (1 mL) was added to a sealable reaction tube. The solution was degassed for 20 minutes and tert-butyl 4-aminobenzoate (**5**) (29.7 mg, 0.154 mmol) was added subsequently. The reaction tube was tightly sealed and the reaction was stirred at 120 °C for 15h. The reaction was cooled to room temperature, diluted with CH_2_Cl_2_, and filtered over a pad of celite. The solvent was removed under reduced pressure. The residue was purified by prep-TLC (Magic base/CH_2_Cl_2_ = 3:1, magic base = CH_2_Cl_2_/MeOH/NH_3_ H_2_O = 60/10/1) to give the desired cpd tert-butyl 4-[2,5-bis[4-(1-piperidyl)piperidine-1-carbonyl]anilino]benzoate (55.5 mg, 0.0844 mmol, yield: 83.7 %). ^1^H NMR (600 MHz, CD_3_OD) δ 7.83 – 7.78 (m, 2H), 7.43 – 7.37 (m, 2H), 7.18 (d, J = 7.5 Hz, 1H), 7.01 (d, J = 8.4 Hz, 2H), 4.66 (t, J = 14.2 Hz, 2H), 3.87 – 3.81 (m, 1H), 3.64 (d, J = 13.3 Hz, 1H), 3.19 – 2.95 (m, 2H), 2.86 – 2.23 (m, 12H), 2.05 – 1.17 (m, 30H). ^13^C NMR (151 MHz, CD_3_OD) δ 170.9, 169.5, 167.3, 149.5, 140.0, 139.1, 132.3, 130.0, 124.3, 122.6, 121.6, 116.0, 81.5, 63.5, 63.3, 51.2, 50.9, 48.5, 48.0, 42.9, 42.6, 29.4, 28.6, 28.6, 28.1, 26.8, 26.7, 25.4. HRMS (ESI) [M+H]^+^ for C_39_H_56_N_5_O_4_: 658.4332, found: 658.4327.

5 mg of the obtained product was dissolved in 1 mL of TFA/CH_2_Cl_2_ (v/v = 1:2). The mixture was stirred at rt for 2h. Concentrated and the crude product (**9**) was used into next step without further purification.

**Biotin-UNC1079\***

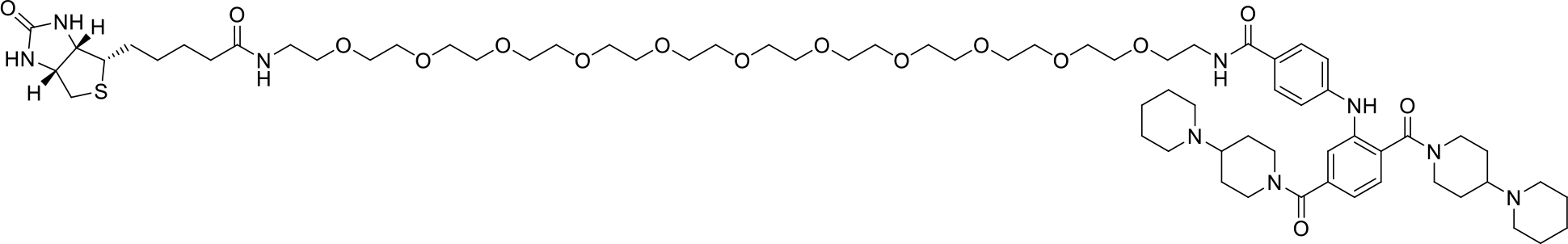

To a mixture of 4-[2,5-bis[4-(1-piperidyl)piperidine-1-carbonyl]anilino]benzoic acid (**9**) (4.50 mg, 0.00748 mmol) and TBTU (2.88 mg, 0.00897 mmol) in DMF (0.3 mL), a solution of Biotin-peg12-NH_2_ (5.77 mg, 0.00748 mmol) and TEA (0.0104 mL, 0.0748 mmol) in DMF (0.2 mL) was added. The mixture was stirred at rt for 12h. The reaction was quenched by the addition of saturated aq. NaHCO_3_ (2 mL) and extracted with CH_2_Cl_2_ (3 mL x 3). The combined organic extracts were dried over Na_2_SO_4_ and concentrated under reduced pressure. The residue was purified by Prep-HPLC (0-70 MeCN/H_2_O with 0.1% FA) to give the desired compound **Biotin-UNC1079\*** (3.60 mg, 0.00266 mmol, yield: 35.5 %). ^1^H NMR (600 MHz, CD_3_OD) δ 8.48 (s, 1H), 7.77 (dd, J = 8.7, 1.6 Hz, 2H), 7.48 – 7.38 (m, 2H), 7.18 (d, J = 7.8 Hz, 1H), 7.08 (dt, J = 9.3, 2.1 Hz, 2H), 4.81 – 4.70 (m, 2H), 4.52 – 4.46 (m, 1H), 4.35 – 4.27 (m, 1H), 4.03 – 3.47 (m, 48H), 3.40 – 3.34 (m, 2H), 3.26 – 2.66 (m, 16H), 2.26 – 2.01 (m, 5H), 1.94 – 1.39 (m, 23H). ^13^C NMR (151 MHz, CD_3_OD) δ 176.1, 171.1, 169.7, 169.5, 166.1, 148.2, 140.9, 139.0, 130.3, 130.2, 127.2, 122.1, 121.2, 117.0, 71.6, 71.3, 71.3, 70.7, 70.6, 64.5, 64.4, 63.4, 61.6, 57.0, 51.3, 51.1, 47.5, 47.0, 42.0, 41.6, 41.1, 40.9, 40.4, 36.7, 29.8, 29.5, 28.3, 27.9, 27.4, 26.9, 24.9, 24.9, 23.3. HRMS (ESI) [M+H]^+^ for C_69_H_112_N_9_O_16_S: 1354.7948, found: 1354.7942.

#### Synthesis of KL-1

**Scheme 4.**
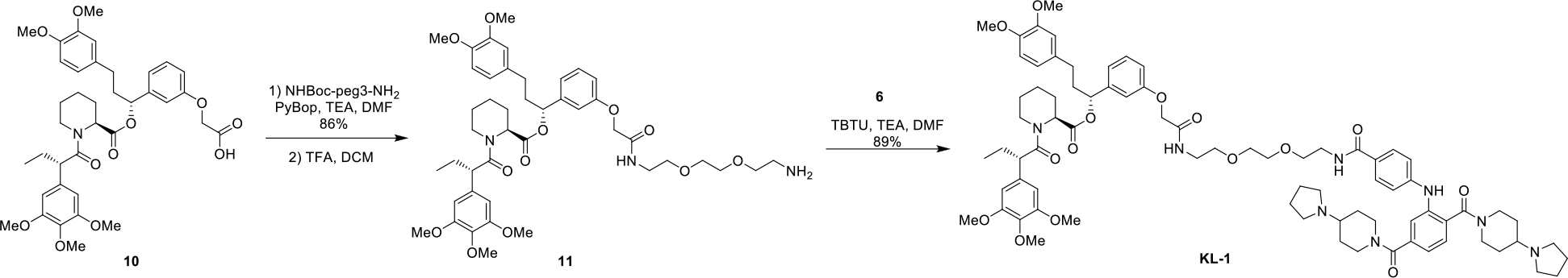
Synthesis of KL-1

**[(1R)-1-[3-[2-[2-[2-(2-aminoethoxy)ethoxy]ethylamino]-2-oxo-ethoxy]phenyl]-3-(3,4-dimethoxyphenyl)propyl](2S)-1-[(2S)-2-(3,4,5-trimethoxyphenyl)butanoyl]piperidine-2-carboxylate (11)**

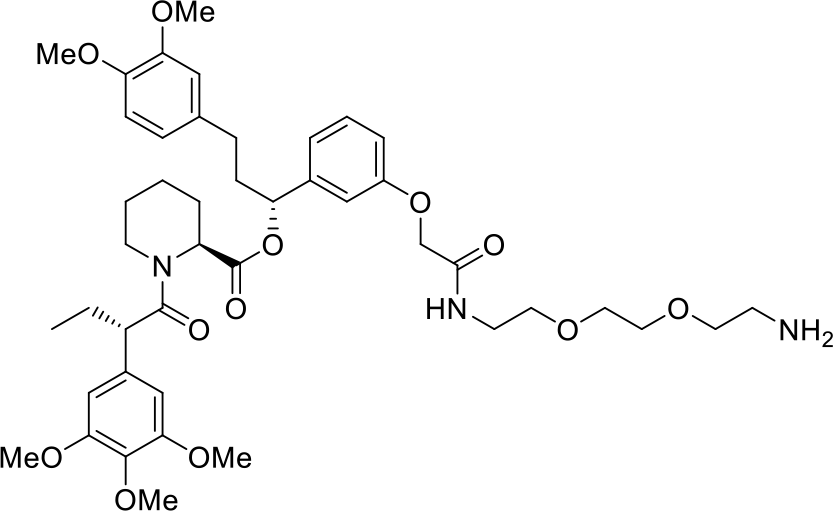

2-[3-[(1R)-3-(3,4-dimethoxyphenyl)-1-[(2S)-1-[(2S)-2-(3,4,5trimethoxyphenyl)butanoyl]piperidine-2-carbonyl]oxy-propyl]phenoxy]acetic acid (**10**) (10.0 mg, 0.0144 mmol), tert-butyl N-[2-[2-(2-aminoethoxy)ethoxy]ethyl]carbamate (4.30 mg, 0.0173 mmol) and TEA (0.00603 mL, 0.0432 mmol) was dissolved in DMF (0.3 mL). PyBOP (9.00 mg, 0.0173 mmol) was added and the reaction was stirred at rt for 12h. The reaction was quenched by the addition of saturated aq. NH_4_Cl (2 mL) and extracted with CH_2_Cl_2_ (3 mL x 3). The combined organic extracts were dried over Na_2_SO_4_ and concentrated under reduced pressure. The residue was purified by Prep-TLC (MeOH/CH_2_Cl_2_ = 5%) to give desired compound [(1R)-1-[3-[2-[2-[2-[2-(tert-butoxycarbonylamino)ethoxy]ethoxy]ethylamino]-2-oxo-ethoxy]phenyl]-3-(3,4-dimethoxyphenyl)propyl](2S)-1-[(2S)-2-(3,4,5-trimethoxyphenyl)butanoyl]piperi-dine-2-carboxylate (11.5 mg, 0.0124 mmol, yield: 86.3 %). ^1^H NMR (600 MHz, CDCl_3_, mixture of rotamers, only major rotamer is reported here) δ 7.21 – 7.15 (m, 1H), 7.11 (s, 1H), 6.81 – 6.75 (m, 3H), 6.68 – 6.61 (m, 3H), 6.41 (s, 2H), 5.62 (dd, J = 8.2, 5.4 Hz, 1H), 5.46 – 5.44 (m, 1H), 4.50 (s, 2H), 3.85 (s, 3H), 3.85 (s, 3H), 3.78 (s, 3H), 3.68 (s, 6H), 3.61 – 3.57 (m, 8H), 3.52 (t, J = 5.3 Hz, 2H), 3.29 (t, J = 5.1 Hz, 2H), 2.79 (td, J = 13.4, 3.0 Hz, 1H), 2.58 – 2.52 (m, 1H), 2.49 – 2.42 (m, 1H), 2.31 (d, J = 13.0 Hz, 1H), 2.11 – 2.02 (m, 3H), 1.96 – 1.88 (m, 2H), 1.75 – 1.67 (m, 3H), 1.60 (d, J = 12.8 Hz, 1H), 1.43 (s, 9H), 1.33 – 1.23 (m, 2H), 0.90 (t, J = 7.3 Hz, 3H). ^13^C NMR (151 MHz, CDCl_3_) δ 172.6, 170.6, 168.3, 157.3, 156.0, 153.5, 153.2, 148.9, 147.3, 142.3, 136.6, 135.3, 133.3, 129.8, 120.2, 119.8, 113.9, 112.8, 111.7, 111.3, 105.0, 75.6, 70.2, 70.2, 70.2, 69.7, 67.3, 60.8, 56.0, 55.9, 55.8, 52.0, 50.8, 43.5, 40.4, 38.8, 38.2, 31.3, 28.4, 28.4, 28.3, 26.8, 25.3, 20.9, 12.5. HRMS (ESI) [M+H]^+^ for C_49_H_70_N_3_O_14_: 924.4858, found: 924.4852.

7.3 mg of the obtained product was dissolved in 1 mL of TFA/CH_2_Cl_2_ (v/v = 1:2). The mixture was stirred at rt for 2h. Concentrated and the crude product (**11**) was used into next step without further purification.

**KL-1**

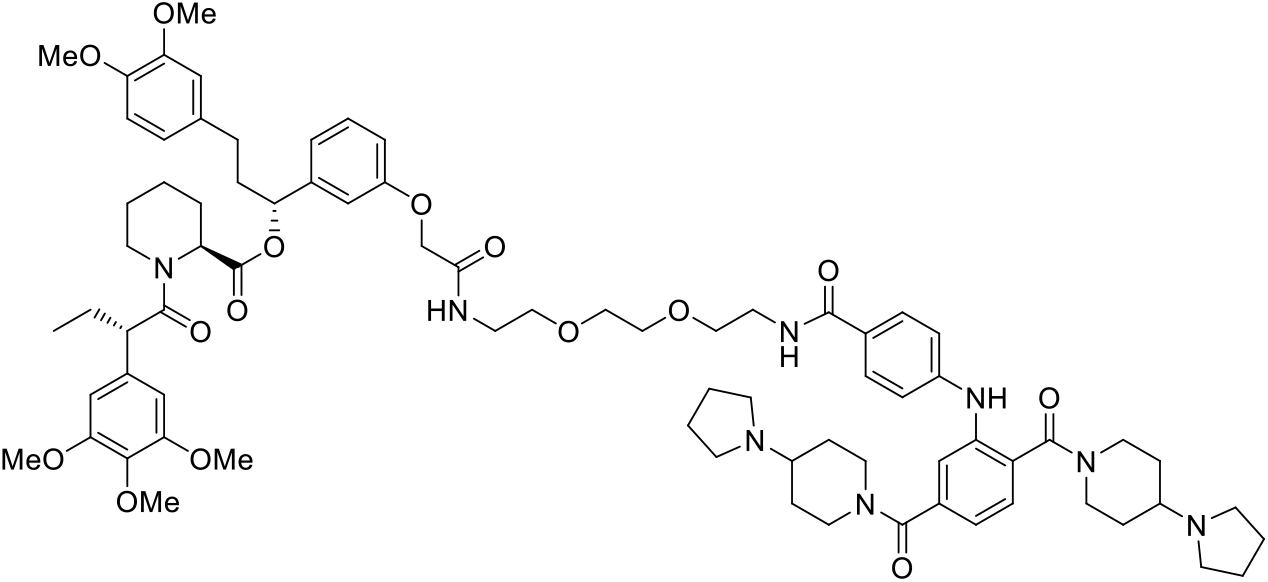

To a mixture of 4-[2,5-bis(4-pyrrolidin-1-ylpiperidine-1-carbonyl)anilino]benzoic acid (**6**) (4.50 mg, 0.00784 mmol) and TBTU (7.56 mg, 0.0235 mmol) in DMF (0.3 mL), a solution of [(1R)-1-[3-[2-[2-[2-(2-aminoethoxy)ethoxy]ethylamino]-2-oxo-ethoxy]phenyl]-3-(3,4-dimethoxyphenyl)propyl] (2S)-1-[(2S)-2-(3,4,5-trimethoxyphenyl)butanoyl]piperidine-2-carboxylate (**11**) (6.46 mg, 0.00784 mmol) and TEA (0.0109 mL, 0.0784 mmol) in DMF (0.3 mL) was added. The mixture was stirred at rt for 12h. The reaction was quenched by the addition of saturated aq. NaHCO_3_ (2 mL) and extracted with CH_2_Cl_2_ (3 mL x 3). The combined organic extracts were dried over Na_2_SO_4_ and concentrated under reduced pressure. The residue was purified by Prep-HPLC (5-90% MeCN/H_2_O with 0.1% FA) to give the desired compound **KL-1** (9.70 mg, 0.00703 mmol, yield: 89.6 %). ^1^H NMR (600 MHz, CD_3_OD, mixture of rotamers, only major rotamer is reported here) δ 7.73 (d, J = 8.2 Hz, 2H), 7.41 (d, J = 7.8 Hz, 1H), 7.36 (s, 1H), 7.18 (t, J = 7.9 Hz, 1H), 7.13 (d, J = 7.8 Hz, 1H), 7.06 (d, J = 8.3 Hz, 2H), 6.84 (d, J = 8.2 Hz, 2H), 6.79 (s, 1H), 6.73 (s, 1H), 6.67 (d, J = 8.2 Hz, 1H), 6.61 – 6.53 (m, 3H), 5.57 (t, J = 6.9 Hz, 1H), 5.38 (d, J = 5.7 Hz, 1H), 4.66 (s, 2H), 4.48 (d, J = 4.5 Hz, 2H), 4.08 (d, J = 13.8 Hz, 1H), 3.87 (t, J = 7.4 Hz, 2H), 3.79 (s, 6H), 3.69 (s, 3H), 3.66 (s, 6H), 3.63 – 3.51 (m, 10H), 3.43 (t, J = 5.8 Hz, 2H), 3.17 – 2.99 (m, 11H), 2.85 (s, 1H), 2.73 (t, J = 13.2 Hz, 1H), 2.58 – 2.51 (m, 1H), 2.47 – 2.38 (m, 1H), 2.28 (d, J = 13.6 Hz, 1H), 2.15 – 1.86 (m, 15H), 1.74 – 1.40 (m, 10H), 1.38 – 1.27 (m, 3H), 0.89 (t, J = 7.3 Hz, 3H). ^13^C NMR (151 MHz, CD_3_OD) δ 175.0, 171.9, 171.2, 171.1, 169.8, 169.6, 159.1, 154.6, 150.4, 148.9, 147.9, 143.7, 141.2, 139.1, 137.9, 137.0, 135.2, 130.9, 130.3, 130.1, 127.4, 121.8, 121.7, 120.7, 119.9, 117.5, 115.1, 114.4, 113.7, 113.3, 106.6, 77.1, 71.3, 71.3, 70.7, 70.4, 68.3, 63.0, 62.9, 61.1, 56.6, 56.6, 56.5, 53.6, 52.7, 52.6, 51.2, 45.0, 40.8, 40.0, 39.3, 32.3, 29.4, 27.6, 26.3, 23.9, 21.9, 12.7. HRMS (ESI) [M+H]^+^ for C_77_H_103_N_8_O_15_: 1379.7543, found: 1379.7537.

#### Synthesis of KL-2

**Scheme 5.**
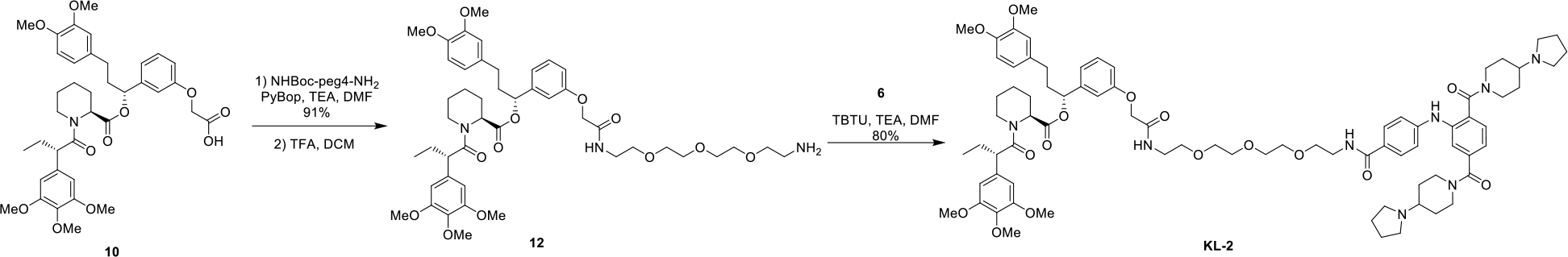
Synthesis of KL-2

**[(1R)-1-[3-[2-[2-[2-[2-(2-aminoethoxy)ethoxy]ethoxy]ethylamino]-2-oxo-ethoxy]phenyl]-3-(3,4-dimethoxyphenyl)propyl] (2S)-1-[(2S)-2-(3,4,5-trimethoxyphenyl)butanoyl]piperidine-2-carboxylate (12)**

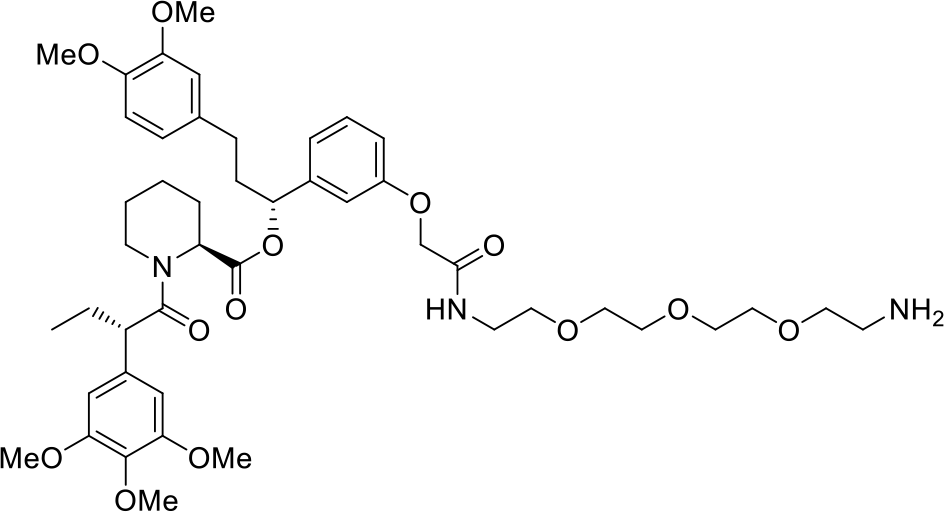

2-[3-[(1R)-3-(3,4-dimethoxyphenyl)-1-[(2S)-1-[(2S)-2-(3,4,5-trimethoxyphenyl)butanoyl]piperidine-2-carbonyl]oxy-propyl]phenoxy]acetic acid (**10**) (10.0 mg, 0.0144 mmol), tert-butyl N-[2-[2-[2-(2-aminoethoxy)ethoxy]ethoxy]ethyl]carbamate (5.06 mg, 0.0173 mmol) and TEA (0.00603 mL, 0.0432 mmol) was dissolved in DMF (0.3 mL). PyBOP (9.00 mg, 0.0173 mmol) was added and the reaction was stirred at rt for 12h. The reaction was quenched by the addition of saturated aq. NH_4_Cl (2 mL) and extracted with CH_2_Cl_2_ (3 mL x 3). The combined organic extracts were dried over Na_2_SO_4_ and concentrated under reduced pressure. The residue was purified by Prep-TLC (MeOH/CH_2_Cl_2_ = 5%) to give desired compound [(1R)-1-[3-[2-[2-[2-[2-[2-(tert-butoxycarbonylamino)ethoxy]ethoxy]ethoxy]ethyl amino]-2-oxo-ethoxy]phenyl]-3-(3,4-dimethoxyphenyl)propyl](2S)-1-[(2S)-2-(3,4,5-trimethoxyphenyl) butanoyl]piperidine-2-carboxylate (12.8 mg, 0.0132 mmol, yield: 91.0 %). ^1^H NMR (600 MHz, CDCl_3_, mixture of rotamers, only major rotamer is reported here) δ 7.18 (td, J = 7.5, 2.0 Hz, 1H), 7.09 (s, 1H), 6.78 (s, 2H), 6.65 (d, J = 11.9 Hz, 3H), 6.41 (s, 2H), 5.62 (dd, J = 8.2, 5.4 Hz, 1H), 5.46 (d, J = 4.7 Hz, 1H), 4.49 (d, J = 1.8 Hz, 2H), 3.84 (s, 3H), 3.84 (s, 3H), 3.78 (s, 3H), 3.69 (s, 6H), 3.64 – 3.58 (m, 12H), 3.51 (t, J = 5.1 Hz, 2H), 3.28 (t, J = 5.1 Hz, 2H), 2.79 (td, J = 13.4, 3.0 Hz, 1H), 2.58 – 2.51 (m, 1H), 2.49 – 2.42 (m, 1H), 2.30 (d, J = 13.4 Hz, 1H), 2.12 – 2.03 (m, 3H), 1.96 – 1.88 (m, 2H), 1.75 – 1.67 (m, 3H), 1.63 – 1.58 (m, 1H), 1.43 (s, 9H), 1.32 – 1.23 (m, 2H), 0.90 (t, J = 7.3 Hz, 3H). ^13^C NMR (151 MHz, CDCl_3_) δ 172.6, 170.6, 168.2, 157.3, 156.0, 153.2, 148.9, 147.3, 142.3, 136.6, 135.3, 133.3, 129.8, 120.2, 119.7, 113.9, 112.8, 111.7, 111.3, 105.0, 75.6, 70.5, 70.5, 70.3, 70.2, 69.7, 67.3, 60.8, 56.0, 55.9, 55.8, 55.8, 52.0, 50.8, 43.5, 38.9, 38.8, 38.3, 31.3, 28.4, 28.4, 28.3, 26.8, 25.3, 20.9, 12.5. HRMS (ESI) [M+H]^+^ for C_51_H_74_N_3_O_15_: 968.5120, found: 968.5115.

7.6 mg of the obtained product was dissolved in 1 mL of TFA/CH_2_Cl_2_ (v/v = 1:2). The mixture was stirred at rt for 2h. Concentrated and the crude product (**12**) was used into next step without further purification.

**KL-2**

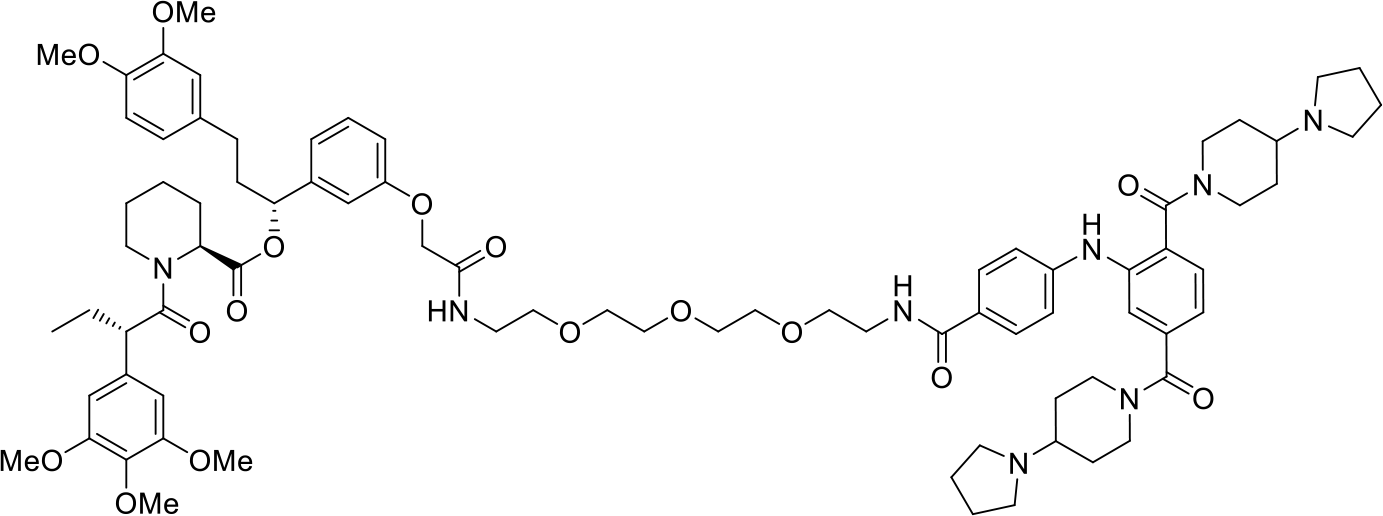

To a mixture of 4-[2,5-bis(4-pyrrolidin-1-ylpiperidine-1-carbonyl)anilino]benzoic acid (**6**) (4.50 mg, 0.00784 mmol) and TBTU (7.56 mg, 0.0235 mmol) in DMF (0.3 mL), a solution of [(1R)-1-[3-[2-[2-[2-[2-(2-aminoethoxy)ethoxy]ethoxy]ethylamino]-2-oxo-ethoxy]phenyl]-3-(3,4-dimethoxyphenyl)propyl] (2S)-1-[(2S)-2-(3,4,5-trimethoxyphenyl)butanoyl]piperidine-2-carboxylate (**12**) (6.81 mg, 0.00784 mmol) and TEA (0.0109 mL, 0.0784 mmol) in DMF (0.3 mL) was added. The mixture was stirred at rt for 12h. The reaction was quenched by the addition of saturated aq. NaHCO_3_ (2 mL) and extracted with CH_2_Cl_2_ (3 mL x 3). The combined organic extracts were dried over Na_2_SO_4_ and concentrated under reduced pressure. The residue was purified by Prep-HPLC (5-90% MeCN/H_2_O with 0.1% FA) to give the desired compound **KL-2** (9.00 mg, 0.00632 mmol, yield: 80.3 %). ^1^H NMR (600 MHz, CD_3_OD, mixture of rotamers, only major rotamer is reported here) δ 7.74 (d, J = 8.2 Hz, 2H), 7.41 (d, J = 7.8 Hz, 1H), 7.37 (s, 1H), 7.18 (t, J = 8.0 Hz, 1H), 7.13 (d, J = 7.8 Hz, 1H), 7.07 (d, J = 8.3 Hz, 2H), 6.85 (d, J = 8.2 Hz, 2H), 6.79 (s, 1H), 6.74 (s, 1H), 6.67 (d, J = 8.1 Hz, 1H), 6.60 – 6.54 (m, 3H), 5.58 (dd, J = 8.4, 5.4 Hz, 1H), 5.38 (d, J = 5.6 Hz, 1H), 4.67 (s, 2H), 4.48 (d, J = 4.9 Hz, 2H), 4.08 (d, J = 13.7 Hz, 1H), 3.87 (t, J = 7.3 Hz, 2H), 3.79 (s, 3H), 3.79 (s, 3H), 3.69 (s, 3H), 3.67 (s, 6H), 3.63 – 3.51 (m, 14H), 3.43 (t, J = 5.8 Hz, 2H), 3.17 – 2.98 (m, 11H), 2.85 (s, 1H), 2.73 (t, J = 13.2 Hz, 1H), 2.58 – 2.51 (m, 1H), 2.46 – 2.39 (m, 1H), 2.28 (d, J = 12.9 Hz, 1H), 2.16 – 1.86 (m, 15H), 1.75 – 1.37 (m, 10H), 1.32 – 1.21 (m, 2H), 0.89 (t, J = 7.3 Hz, 3H). ^13^C NMR (151 MHz, CD_3_OD) δ 175.0, 171.9, 171.2, 171.0, 169.8, 169.6, 159.1, 154.6, 150.4, 148.9, 147.9, 143.7, 141.3, 139.1, 137.9, 137.0, 135.2, 131.0, 130.3, 130.1, 127.5, 121.8, 121.7, 121.7, 120.6, 119.9, 117.5, 115.1, 114.4, 113.7, 113.3, 106.6, 77.1, 71.6, 71.3, 70.7, 70.4, 68.3, 63.0, 62.9, 61.1, 56.6, 56.5, 53.6, 52.7, 52.6, 51.2, 45.0, 40.9, 40.0, 39.3, 32.3, 29.4, 27.6, 26.3, 23.9, 21.9, 12.7. HRMS (ESI) [M+H]^+^ for C_79_H_107_N_8_O_16_: 1423.7805, found: 1423.7800.

#### Synthesis of KL-3

**Scheme 6.**
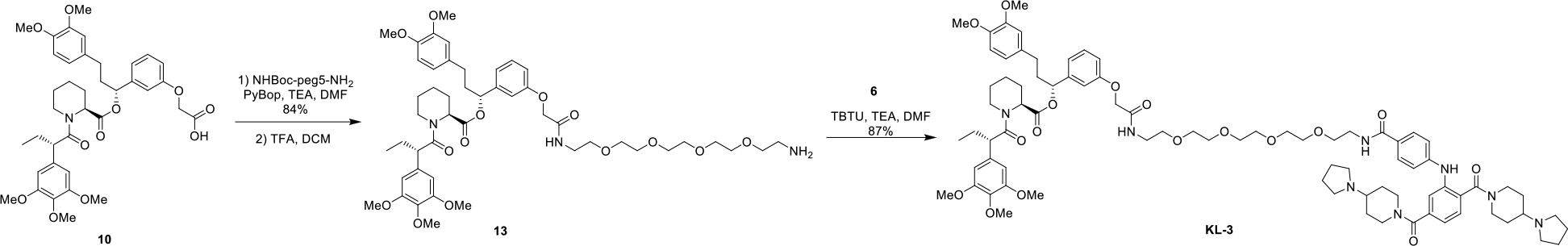
Synthesis of KL-3

**[(1R)-1-[3-[2-[2-[2-[2-[2-(2-aminoethoxy)ethoxy]ethoxy]ethoxy]ethylamino]-2-oxo-ethoxy]phenyl]-3-(3,4-dimethoxyphenyl)propyl](2S)-1-[(2S)-2-(3,4,5-trimethoxyphenyl) butanoyl]piperidine-2-carboxylate (13)**

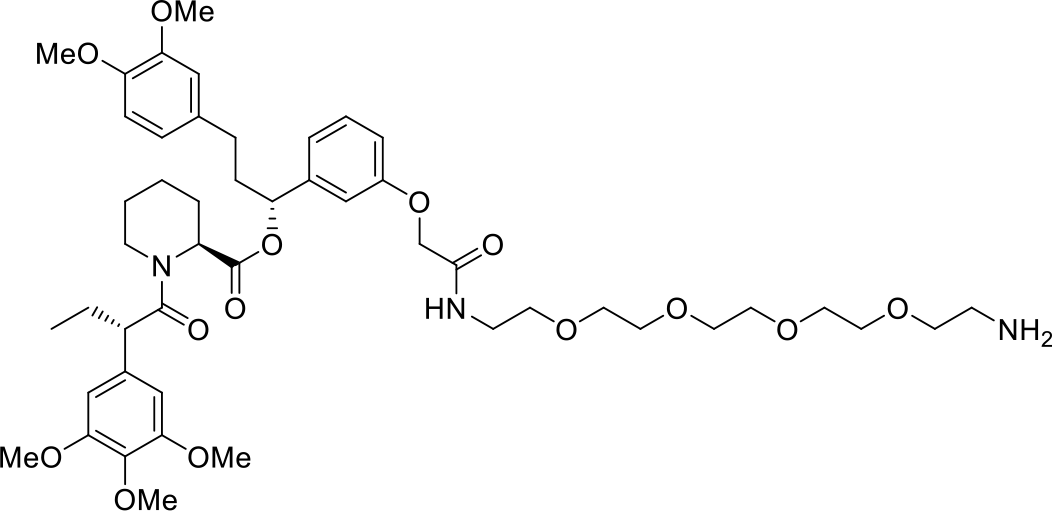

2-[3-[(1R)-3-(3,4-dimethoxyphenyl)-1-[(2S)-1-[(2S)-2-(3,4,5-trimethoxyphenyl)butanoyl]piperidine-2-carbonyl]oxy-propyl]phenoxy]acetic acid (**10**) (10.0 mg, 0.0144 mmol), tert-butyl N-[2-[2-[2-[2-(2-aminoethoxy)ethoxy]ethoxy]ethoxy]ethyl]carbamate (5.82 mg, 0.0173 mmol) and TEA (0.00603 mL, 0.0432 mmol) was dissolved in DMF (0.3 mL). PyBOP (9.00 mg, 0.0173 mmol) was added and the reaction was stirred at rt for 12h. The reaction was quenched by the addition of saturated aq. NH_4_Cl (2 mL) and extracted with CH_2_Cl_2_ (3 mL x 3). The combined organic extracts were dried over Na_2_SO_4_ and concentrated under reduced pressure. The residue was purified by Prep-TLC (MeOH/CH_2_Cl_2_ = 5%) to give desired compound [(1R)-1-[3-[2-[2-[2-[2-[2-[2-(tert-butoxycarbonylamino)ethoxy]ethoxy]ethoxy]ethoxy] ethylamino]-2-oxo-ethoxy]phenyl]-3-(3,4-dimethoxyphenyl)propyl](2S)-1-[(2S)-2-(3,4,5-trimethoxy phenyl)butanoyl]piperidine-2-carboxylate (12.2 mg, 0.0121 mmol, yield: 83.6 %). ^1^H NMR (600 MHz, CDCl_3_, mixture of rotamers, only major rotamer is reported here) δ 7.18 (t, J = 7.8 Hz, 1H), 7.13 (s, 1H), 6.78 (d, J = 4.1 Hz, 2H), 6.67 – 6.61 (m, 3H), 6.41 (s, 2H), 5.62 (dd, J = 8.2, 5.5 Hz, 1H), 5.49 – 5.44 (m, 1H), 4.49 (s, 2H), 3.85 (s, 3H), 3.84 (s, 3H), 3.78 (s, 3H), 3.69 (s, 6H), 3.65 – 3.58 (m, 16H), 3.51 (t, J = 5.1 Hz, 2H), 3.29 (t, J = 5.0 Hz, 2H), 2.79 (td, J = 13.4, 3.0 Hz, 1H), 2.57 – 2.51 (m, 1H), 2.49 – 2.42 (m, 1H), 2.30 (d, J = 13.4 Hz, 1H), 2.12 – 2.02 (m, 4H), 1.96 – 1.88 (m, 1H), 1.75 – 1.67 (m, 3H), 1.60 (d, J = 13.8 Hz, 1H), 1.43 (s, 9H), 1.31 – 1.21 (m, 2H), 0.90 (t, J = 7.2 Hz, 3H). ^13^C NMR (151 MHz, CDCl_3_) δ 172.6, 170.5, 168.2, 157.3, 156.0, 153.5, 153.2, 148.9, 147.3, 142.3, 136.6, 135.3, 133.3, 129.8, 120.2, 119.7, 114.0, 112.8, 111.7, 111.3, 105.0, 75.6, 70.5, 70.5, 70.3, 70.2, 69.7, 67.3, 60.8, 56.3, 56.0, 55.9, 55.9, 55.8, 52.0, 50.8, 43.5, 38.8, 38.2, 31.3, 28.4, 28.4, 28.3, 26.8, 25.3, 20.9, 12.5. HRMS (ESI) [M+H]^+^ for C_53_H_78_N_3_O_16_: 1012.5382, found: 1012.5377.

8.0 mg of the obtained product was dissolved in 1 mL of TFA/CH_2_Cl_2_ (v/v = 1:2). The mixture was stirred at rt for 2h. Concentrated and the crude product (**13**) was used into next step without further purification.

**KL-3**

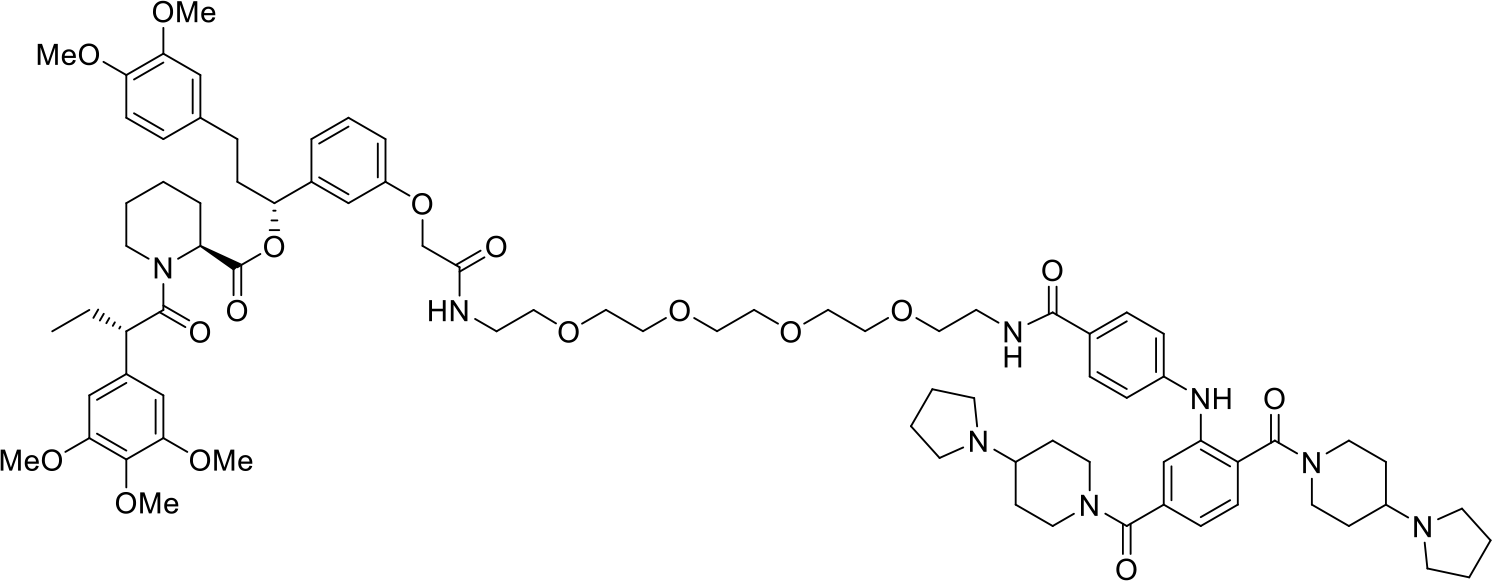

To a mixture of 4-[2,5-bis(4-pyrrolidin-1-ylpiperidine-1-carbonyl)anilino]benzoic acid (**6**) (4.50 mg, 0.00784 mmol) and TBTU (7.56 mg, 0.0235 mmol) in DMF (0.3 mL), a solution of [(1R)-1-[3-[2-[2-[2-[2-[2-(2-aminoethoxy)ethoxy]ethoxy]ethoxy]ethylamino]-2-oxo-ethoxy]phenyl]-3-(3,4-dimethoxy phenyl)propyl](2S)-1-[(2S)-2-(3,4,5-trimethoxyphenyl)butanoyl]piperidine-2-carboxylate (**13**) (7.15 mg, 0.00784 mmol) and TEA (0.0109 mL, 0.0784 mmol) in DMF (0.3 mL) was added. The mixture was stirred at rt for 12h. The reaction was quenched by the addition of saturated aq. NaHCO_3_ (2 mL) and extracted with CH_2_Cl_2_ (3 mL x 3). The combined organic extracts were dried over Na_2_SO_4_ and concentrated under reduced pressure. The residue was purified by Prep-HPLC (5-90% MeCN/H_2_O with 0.1% FA) to give the desired compound **KL-3** (10.1 mg, 0.00688 mmol, yield: 87.3 %). ^1^H NMR (600 MHz, CD_3_OD, mixture of rotamers, only major rotamer is reported here) δ 7.75 (d, J = 8.2 Hz, 2H), 7.41 (d, J = 7.8 Hz, 1H), 7.37 (s, 1H), 7.18 (t, J = 7.9 Hz, 1H), 7.13 (d, J = 7.8 Hz, 1H), 7.07 (d, J = 8.3 Hz, 2H), 6.85 (t, J = 7.3 Hz, 2H), 6.80 (s, 1H), 6.74 (s, 1H), 6.67 (d, J = 8.2 Hz, 1H), 6.61 – 6.53 (m, 3H), 5.63 – 5.51 (m, 1H), 5.38 (d, J = 5.6 Hz, 1H), 4.68 (s, 2H), 4.50 (d, J = 4.6 Hz, 2H), 4.08 (d, J = 13.7 Hz, 1H), 3.87 (t, J = 7.3 Hz, 2H), 3.80 (s, 3H), 3.79 (s, 3H), 3.69 (s, 3H), 3.67 (s, 6H), 3.63 – 3.52 (m, 18H), 3.44 (t, J = 5.9 Hz, 2H), 3.19 – 2.99 (m, 11H), 2.86 (s, 1H), 2.73 (t, J = 13.1 Hz, 1H), 2.58 – 2.50 (m, 1H), 2.47 – 2.40 (m, 1H), 2.29 (d, J = 12.8 Hz, 1H), 2.15 – 1.87 (m, 15H), 1.75 – 1.39 (m, 10H), 1.34 – 1.21 (m, 2H), 0.89 (t, J = 7.2 Hz, 3H). ^13^C NMR (151 MHz, CD_3_OD) δ 175.0, 171.9, 171.2, 171.0, 169.8, 169.6, 159.1, 154.6, 150.5, 148.9, 147.9, 143.7, 141.3, 139.0, 137.9, 137.0, 135.2, 131.0, 130.3, 130.1, 127.5, 121.8, 121.7, 120.7, 119.9, 117.5, 115.1, 114.4, 113.7, 113.3, 106.6, 77.1, 71.6, 71.5, 71.3, 71.3, 70.7, 70.4, 68.3, 63.0, 62.9, 61.1, 56.6, 56.5, 53.6, 52.7, 52.6, 51.2, 45.0, 40.9, 40.0, 39.3, 32.3, 29.4, 27.6, 26.3, 23.9, 21.9, 12.7. HRMS (ESI) [M+H]^+^ for C_81_H_111_N_8_O_17_: 1467.8067, found: 1467.8062.

#### Synthesis of KL-4

**Scheme 7.**
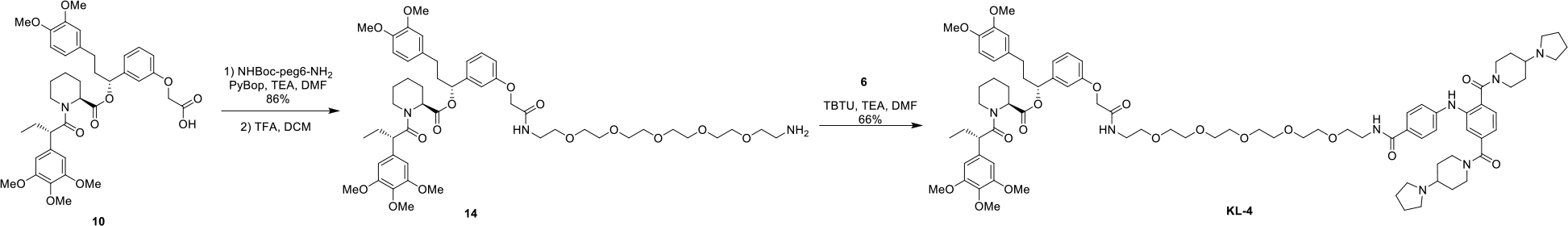
Synthesis of KL-4

**[(1R)-1-[3-[2-[2-[2-[2-[2-[2-(2-aminoethoxy)ethoxy]ethoxy]ethoxy]ethoxy]ethylamino]-2-oxo-ethoxy]phenyl]-3-(3,4-dimethoxyphenyl)propyl](2S)-1-[(2S)-2-(3,4,5-trimethoxyphenyl) butanoyl]piperidine-2-carboxylate (14)**

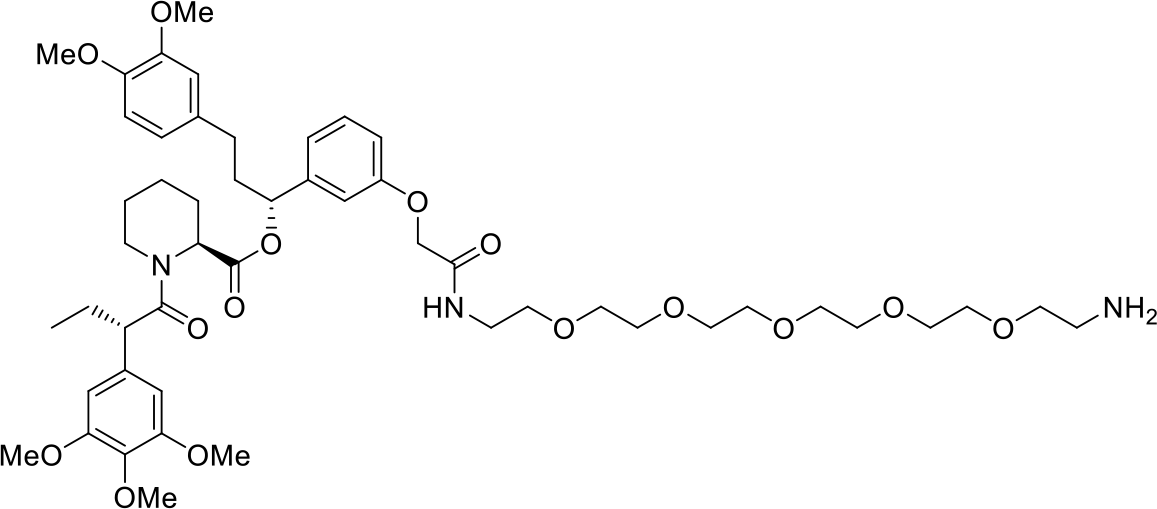

2-[3-[(1R)-3-(3,4-dimethoxyphenyl)-1-[(2S)-1-[(2S)-2-(3,4,5-trimethoxyphenyl)butanoyl]piperidine-2-carbonyl]oxy-propyl]phenoxy]acetic acid (**10**) (10.0 mg, 0.0144 mmol), tert-butyl N-[2-[2-[2-[2-[2-(2-aminoethoxy)ethoxy]ethoxy]ethoxy]ethoxy]ethyl]carbamate (6.58 mg, 0.0173 mmol) and TEA (0.00603 mL, 0.0432 mmol) was dissolved in DMF (0.3 mL). PyBOP (9.00 mg, 0.0173 mmol) was added and the reaction was stirred at rt for 12h. The reaction was quenched by the addition of saturated aq. NH_4_Cl (2 mL) and extracted with CH_2_Cl_2_ (3 mL x 3). The combined organic extracts were dried over Na_2_SO_4_ and concentrated under reduced pressure. The residue was purified by Prep-TLC (MeOH/CH_2_Cl_2_ = 5%) to give desired compound [(1R)-1-[3-[2-[2-[2-[2-[2-[2-[2-(tert-butoxycarbonylamino)ethoxy]ethoxy]ethoxy] ethoxy]ethoxy]ethylamino]-2-oxo-ethoxy]phenyl]-3-(3,4-dimethoxyphenyl)propyl] (2S)-1-[(2S)-2-(3,4,5-trimethoxyphenyl)butanoyl]piperidine-2-carboxylate (13.2 mg, 0.0125 mmol, yield: 86.7 %). ^1^H NMR (600 MHz, CDCl_3_, mixture of rotamers, only major rotamer is reported here) δ 7.14 (t, J = 7.8 Hz, 1H), 7.04 (t, J = 5.9 Hz, 1H), 6.74 (s, 2H), 6.61 (d, J = 8.7 Hz, 3H), 6.38 (s, 2H), 5.59 (dd, J = 8.2, 5.4 Hz, 1H), 5.46 – 5.40 (m, 1H), 5.06 (s, 1H), 4.45 (s, 2H), 3.81 (s, 3H), 3.80 (s, 3H), 3.74 (s, 3H), 3.65 (s, 6H), 3.60 – 3.55 (m, 20H), 3.48 (t, J = 5.0 Hz, 2H), 3.25 (s, 2H), 2.75 (td, J = 13.3, 3.0 Hz, 1H), 2.54 – 2.47 (m, 1H), 2.46 – 2.38 (m, 1H), 2.27 (d, J = 14.6 Hz, 1H), 2.08 – 2.00 (m, 2H), 1.94 – 1.84 (m, 1H), 1.71 – 1.63 (m, 3H), 1.57 (d, J = 13.4 Hz, 1H), 1.39 (s, 9H), 1.28 – 1.17 (m, 2H), 0.86 (t, J = 7.3 Hz, 3H). ^13^C NMR (151 MHz, CDCl_3_) δ 172.6, 170.5, 168.1, 157.3, 153.1, 148.8, 147.3, 142.2, 136.6, 135.3, 133.3, 129.8, 120.1, 119.7, 114.0, 112.8, 111.7, 111.3, 104.9, 75.6, 70.5, 70.4, 70.2, 70.2, 70.2, 69.7, 67.3, 60.7, 56.2, 55.9, 55.9, 55.8, 55.8, 52.0, 50.7, 43.4, 40.3, 38.8, 38.8, 38.2, 31.2, 28.4, 28.3, 26.7, 25.3, 20.9, 12.5. HRMS (ESI) [M+H]^+^ for C_55_H_82_N_3_O_17_: 1056.5644, found: 1056.5639.

8.3 mg of the obtained product was dissolved in 1 mL of TFA/CH_2_Cl_2_ (v/v = 1:2). The mixture was stirred at rt for 2h. Concentrated and the crude product (**14**) was used into next step without further purification.

**KL-4**

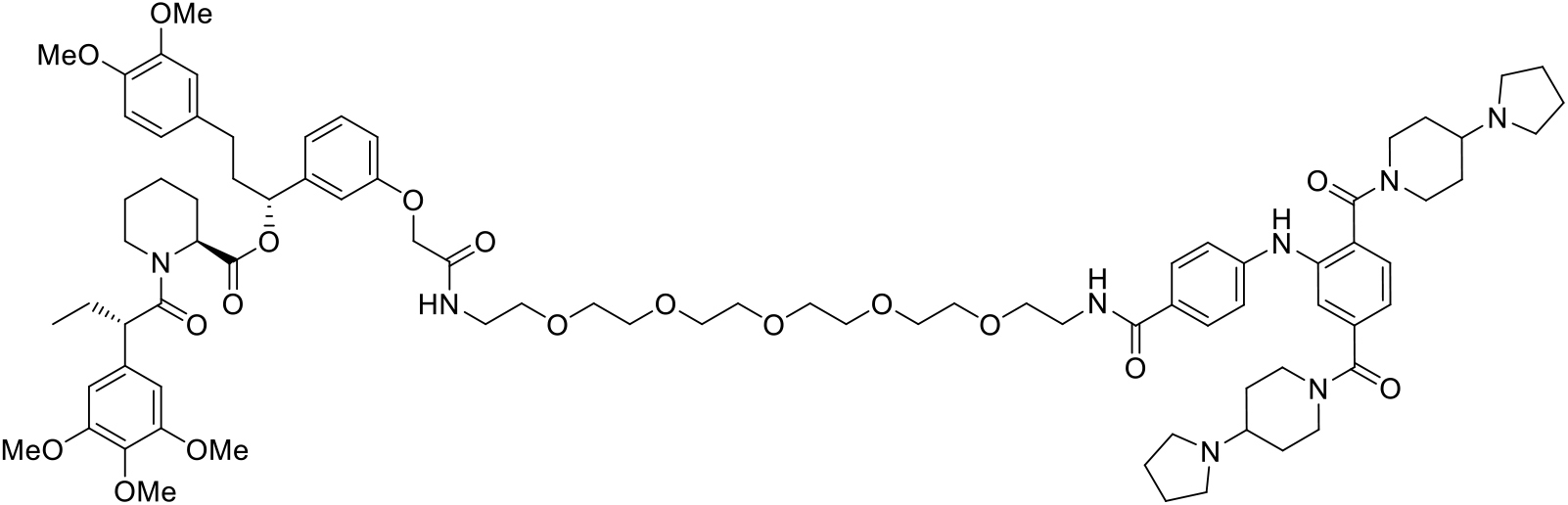

To a mixture of 4-[2,5-bis(4-pyrrolidin-1-ylpiperidine-1-carbonyl)anilino]benzoic acid (**6**) (4.50 mg, 0.00784 mmol) and TBTU (7.56 mg, 0.0235 mmol) in DMF (0.3 mL), a solution of [(1R)-1-[3-[2-[2-[2-[2-[2-[2-(2-aminoethoxy)ethoxy]ethoxy]ethoxy]ethoxy]ethylamino]-2-oxo-ethoxy]phenyl]-3-(3,4-dimethoxyphenyl)propyl] (2S)-1-[(2S)-2-(3,4,5-trimethoxyphenyl)butanoyl]piperidine-2-carboxylate (**14**) (7.50 mg, 0.00784 mmol) and TEA (0.0109 mL, 0.0784 mmol) in DMF (0.3 mL) was added. The mixture was stirred at rt for 12h. The reaction was quenched by the addition of saturated aq. NaHCO_3_ (2 mL) and extracted with CH_2_Cl_2_ (3 mL x 3). The combined organic extracts were dried over Na_2_SO_4_ and concentrated under reduced pressure. The residue was purified by Prep-HPLC (5-90% MeCN/H_2_O with 0.1% FA) to give the desired compound **KL-4** (7.90 mg, 0.00523 mmol, yield: 66.6 %). ^1^H NMR (600 MHz, CD_3_OD, mixture of rotamers, only major rotamer is reported here) δ 7.75 (d, J = 8.3 Hz, 2H), 7.40 (d, J = 7.9 Hz, 1H), 7.36 (s, 1H), 7.19 (t, J = 8.0 Hz, 1H), 7.12 (d, J = 7.8 Hz, 1H), 7.07 (d, J = 8.3 Hz, 2H), 6.86 (t, J = 8.1 Hz, 2H), 6.80 (s, 1H), 6.74 (s, 1H), 6.68 (d, J = 8.2 Hz, 1H), 6.61 – 6.53 (m, 3H), 5.58 (dd, J = 8.3, 5.5 Hz, 1H), 5.39 (d, J = 5.6 Hz, 1H), 4.65 (s, 2H), 4.51 (d, J = 4.3 Hz, 2H), 4.08 (d, J = 13.7 Hz, 1H), 3.87 (t, J = 7.3 Hz, 2H), 3.80 (s, 3H), 3.79 (s, 3H), 3.69 (s, 3H), 3.67 (s, 6H), 3.63 – 3.53 (m, 22H), 3.45 (d, J = 5.6 Hz, 2H), 3.08 – 2.84 (m, 12H), 2.73 (td, J = 13.5, 3.0 Hz, 2H), 2.57 – 2.51 (m, 1H), 2.47 – 2.40 (m, 1H), 2.29 (d, J = 13.6 Hz, 1H), 2.08 – 1.86 (m, 15H), 1.75 – 1.42 (m, 10H), 1.33 – 1.19 (m, 2H), 0.89 (t, J = 7.3 Hz, 3H). ^13^C NMR (151 MHz, CD_3_OD) δ 175.0, 171.9, 171.1, 171.0, 169.7, 169.6, 159.1, 154.6, 150.4, 148.9, 147.9, 143.7, 141.2, 139.1, 137.9, 137.0, 135.1, 130.9, 130.3, 130.1, 127.4, 121.8, 121.7, 120.6, 119.9, 117.5, 115.1, 114.4, 113.7, 113.3, 106.6, 77.1, 71.6, 71.5, 71.5, 71.3, 70.7, 70.4, 68.3, 63.0, 62.9, 61.1, 56.6, 56.6, 56.5, 53.6, 52.7, 52.5, 51.2, 45.0, 40.9, 40.0, 39.3, 32.3, 29.4, 27.6, 26.3, 23.9, 21.9, 12.7. HRMS (ESI) [M+H]^+^ for C_83_H_115_N_8_O_18_: 1511.8329, found: 1511.8324.

#### Synthesis of KL-5

**Scheme 8.**
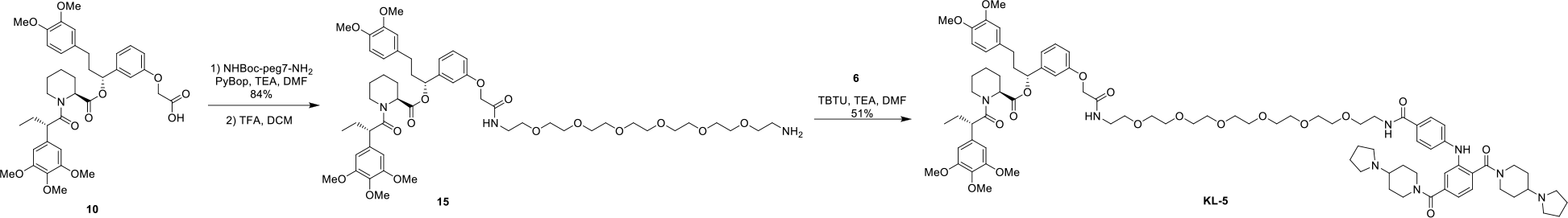
Synthesis of KL-5

**[(1R)-1-[3-[2-[2-[2-[2-[2-[2-[2-(2-aminoethoxy)ethoxy]ethoxy]ethoxy]ethoxy]ethoxy]ethylamino]-2-oxo-ethoxy]phenyl]-3-(3,4-dimethoxyphenyl)propyl] (2S)-1-[(2S)-2-(3,4,5-trimethoxyphenyl) butanoyl]piperidine-2-carboxylate (15)**

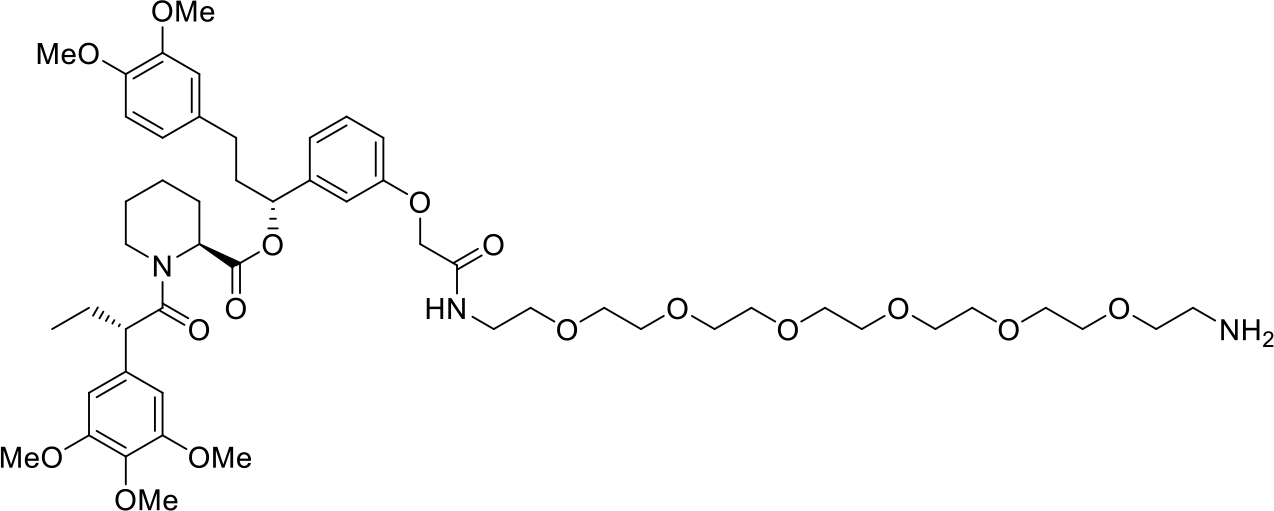

2-[3-[(1R)-3-(3,4-dimethoxyphenyl)-1-[(2S)-1-[(2S)-2-(3,4,5-trimethoxyphenyl)butanoyl]piperidine-2-carbonyl]oxy-propyl]phenoxy]acetic acid (**10**) (10.0 mg, 0.0144 mmol), tert-butyl N-[2-[2-[2-[2-[2-[2-(2-aminoethoxy)ethoxy]ethoxy]ethoxy]ethoxy]ethoxy]ethyl]carbamate (7.34 mg, 0.0173 mmol) and TEA (0.00603 mL, 0.0432 mmol) was dissolved in DMF (0.3 mL). PyBOP (9.00 mg, 0.0173 mmol) was added and the reaction was stirred at rt for 12h. The reaction was quenched by the addition of saturated aq. NH_4_Cl (2 mL) and extracted with CH_2_Cl_2_ (3 mL x 3). The combined organic extracts were dried over Na_2_SO_4_ and concentrated under reduced pressure. The residue was purified by Prep-TLC (MeOH/CH_2_Cl_2_ = 5%) to give desired compound [(1R)-1-[3-[2-[2-[2-[2-[2-[2-[2-[2-(tert-butoxycarbonylamino)ethoxy] ethoxy]ethoxy]ethoxy]ethoxy]ethoxy]ethylamino]-2-oxo-ethoxy]phenyl]-3-(3,4-dimethoxyphenyl)propyl] (2S)-1-[(2S)-2-(3,4,5-trimethoxyphenyl)butanoyl]piperidine-2-carboxylate (13.4 mg, 0.0122 mmol, yield: 84.5 %). ^1^H NMR (500 MHz, CDCl_3_, mixture of rotamers, only major rotamer is reported here) δ 7.18 (t, J = 7.8 Hz, 1H), 7.08 (d, J = 5.6 Hz, 1H), 6.78 (d, J = 1.3 Hz, 2H), 6.68 – 6.62 (m, 3H), 6.41 (s, 2H), 5.63 (dd, J = 8.2, 5.4 Hz, 1H), 5.46 (d, J = 4.6 Hz, 1H), 4.49 (s, 2H), 3.85 (s, 3H), 3.85 (s, 3H), 3.78 (s, 3H), 3.69 (s, 6H), 3.63 (dd, J = 8.2, 3.1 Hz, 20H), 3.53 (t, J = 5.2 Hz, 2H), 3.30 (t, J = 5.2 Hz, 2H), 2.80 (td, J = 13.3, 3.0 Hz, 1H), 2.58 – 2.51 (m, 1H), 2.50 – 2.43 (m, 1H), 2.30 (d, J = 13.4 Hz, 1H), 2.12 – 2.03 (m, 3H), 1.97 – 1.89 (m, 1H), 1.75 – 1.66 (m, 3H), 1.61 (d, J = 13.7 Hz, 1H), 1.44 (s, 9H), 1.31 – 1.22 (m, 2H), 0.90 (t, J = 7.3 Hz, 3H). ^13^C NMR (126 MHz, CDCl_3_) δ 172.6, 170.6, 168.2, 157.3, 156.0, 153.2, 148.9, 147.4, 142.3, 136.7, 135.3, 133.4, 129.8, 120.2, 119.7, 114.0, 112.8, 111.7, 111.3, 105.0, 75.7, 70.5, 70.5, 70.3, 70.2, 69.7, 67.3, 60.8, 56.0, 55.9, 55.8, 52.1, 50.8, 43.5, 40.5, 38.8, 38.3, 31.3, 28.4, 26.8, 25.3, 20.9, 12.5. HRMS (ESI) [M+H]^+^ for C_57_H_86_N_3_O_18_: 1100.5906, found: 1100.5901.

8.6 mg of the obtained product was dissolved in 1 mL of TFA/CH_2_Cl_2_ (v/v = 1:2). The mixture was stirred at rt for 2h. Concentrated and the crude product (**15**) was used into next step without further purification.

**KL-5**

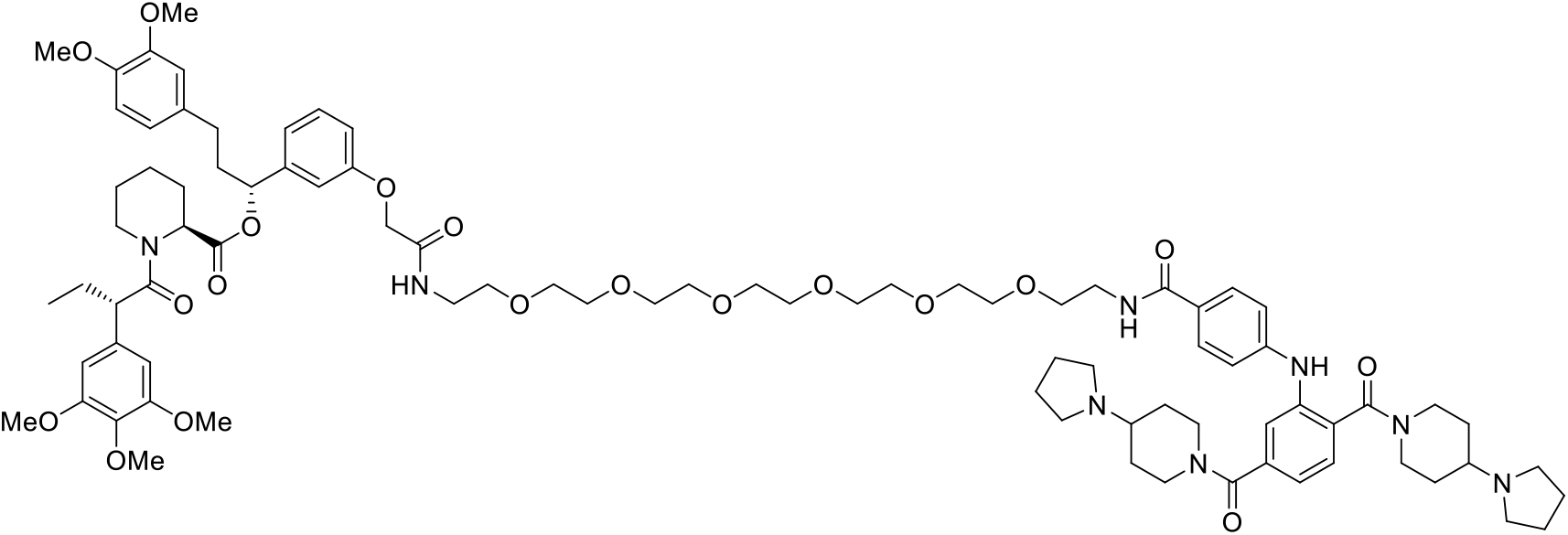

To a mixture of 4-[2,5-bis(4-pyrrolidin-1-ylpiperidine-1-carbonyl)anilino]benzoic acid (**6**) (4.50 mg, 0.00784 mmol) and TBTU (7.56 mg, 0.0235 mmol) in DMF (0.3 mL), a solution of [(1R)-1-[3-[2-[2-[2-[2-[2-[2-[2-(2-aminoethoxy)ethoxy]ethoxy]ethoxy]ethoxy]ethoxy]ethylamino]-2-oxo-ethoxy]phenyl]-3-(3,4-dimethoxyphenyl)propyl] (2S)-1-[(2S)-2-(3,4,5-trimethoxyphenyl)butanoyl]piperidine-2-carboxy-late (**15**) (7.84 mg, 0.00784 mmol) and TEA (0.0109 mL, 0.0784 mmol) in DMF (0.3 mL) was added. The mixture was stirred at rt for 12h. The reaction was quenched by the addition of saturated aq. NaHCO_3_ (2 mL) and extracted with CH_2_Cl_2_ (3 mL x 3). The combined organic extracts were dried over Na_2_SO_4_ and concentrated under reduced pressure. The residue was purified by Prep-HPLC (5-90% MeCN/H_2_O with 0.1% FA) to give the desired compound **KL-5** (6.20 mg, 0.00398 mmol, yield: 50.8 %). ^1^H NMR (600 MHz, CD_3_OD, mixture of rotamers, only major rotamer is reported here) δ 7.75 (d, J = 8.3 Hz, 2H), 7.41 (d, J = 7.8 Hz, 1H), 7.36 (s, 1H), 7.19 (t, J = 8.0 Hz, 1H), 7.12 (d, J = 7.8 Hz, 1H), 7.08 (d, J = 8.3 Hz, 2H), 6.86 (t, J = 8.1 Hz, 2H), 6.80 (s, 1H), 6.74 (s, 1H), 6.68 (d, J = 8.2 Hz, 1H), 6.61 – 6.53 (m, 3H), 5.59 (dd, J = 8.3, 5.5 Hz, 1H), 5.39 (d, J = 5.6 Hz, 1H), 4.64 (s, 2H), 4.51 (d, J = 4.3 Hz, 2H), 4.08 (d, J = 13.7 Hz, 1H), 3.87 (t, J = 7.3 Hz, 2H), 3.80 (s, 3H), 3.79 (s, 3H), 3.69 (s, 3H), 3.67 (s, 6H), 3.63 – 3.54 (m, 26H), 3.46 (d, J = 5.6 Hz, 2H), 3.04 – 2.81 (m, 11H), 2.76 – 2.70 (m, 2H), 2.58 – 2.51 (m, 1H), 2.47 – 2.39 (m, 1H), 2.29 (d, J = 13.4 Hz, 1H), 2.07 – 1.85 (m, 15H), 1.75 – 1.37 (m, 10H), 1.33 – 1.15 (m, 2H), 0.89 (t, J = 7.3 Hz, 3H). ^13^C NMR (151 MHz, CD_3_OD) δ 175.0, 171.9, 171.1, 171.0, 169.7, 169.6, 159.2, 154.6, 150.5, 148.9, 147.9, 143.7, 141.2, 139.1, 137.9, 137.0, 135.2, 130.9, 130.3, 130.1, 127.4, 121.8, 121.6, 120.7, 119.9, 117.5, 115.2, 114.4, 113.7, 113.3, 106.6, 77.1, 71.6, 71.5, 71.5, 71.3, 70.7, 70.4, 68.3, 63.0, 62.9, 61.1, 56.6, 56.6, 56.5, 53.6, 52.7, 52.5, 51.2, 45.0, 40.9, 40.0, 39.3, 32.3, 29.4, 27.6, 26.3, 24.0, 21.9, 12.7. HRMS (ESI) [M+H]^+^ for C_85_H_119_N_8_O_19_: 1555.8591, found: 1555.8586.

#### Synthesis of KL-6

**Scheme 9.**
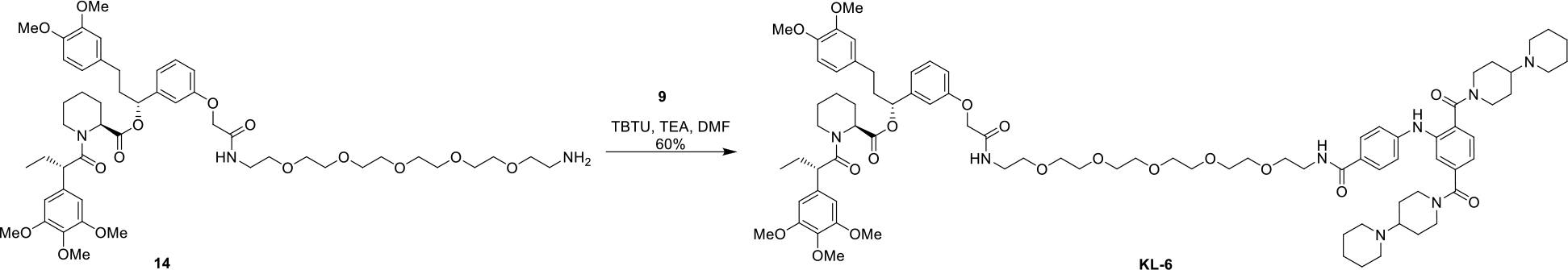
Synthesis of KL-6

**KL-6**

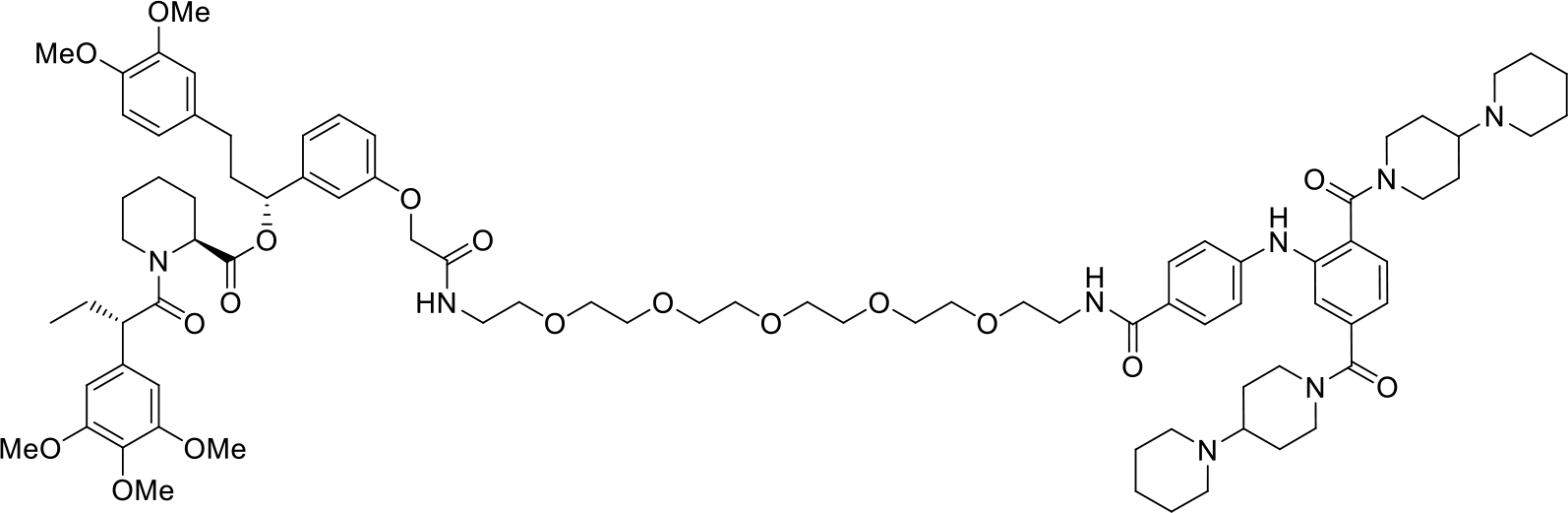

To a mixture of 4-[2,5-bis[4-(1-piperidyl)piperidine-1-carbonyl]anilino]benzoic acid (**9**) (9.50 mg, 0.0158 mmol) and TBTU (7.56 mg, 0.0235 mmol) in DMF (0.3 mL), a solution of N-[2-[2-[2-[2-[2-(2-aminoethoxy)ethoxy]ethoxy]ethoxy]ethoxy]ethyl]-2-[(9S)-7-(4-chlorophenyl)-4,5,13-trimethyl-3-thia-1,8,11,12-tetrazatricyclo[8.3.0.02,6]trideca-2(6),4,7,10,12-pentaen-9-yl]acetamide (**14**) (15.6 mg, 0.0235 mmol) and TEA (0.0219 mL, 0.157 mmol) in DMF (0.3 mL) was added. The mixture was stirred at rt for 12h. The reaction was quenched by the addition of saturated aq. NaHCO_3_ (2 mL) and extracted with CH_2_Cl_2_ (3 mL x 3). The combined organic extracts were dried over Na_2_SO_4_ and concentrated under reduced pressure. The residue was purified by Prep-HPLC (5-90% MeCN/H_2_O with 0.1%FA) to give the desired compound **KL-6** (14.6 mg, 0.00948 mmol, yield: 60.4 %). ^1^H NMR (600 MHz, CD_3_OD, mixture of rotamers, only major rotamer is reported here) δ 8.51 (s, 1H), 7.75 (d, J = 8.3 Hz, 2H), 7.46 – 7.38 (m, 2H), 7.21 – 7.14 (m, 2H), 7.06 (d, J = 8.3 Hz, 2H), 6.86 (td, J = 8.0, 1.9 Hz, 2H), 6.80 (s, 1H), 6.74 (s, 1H), 6.68 (d, J = 8.2 Hz, 1H), 6.62 – 6.53 (m, 3H), 5.58 (dd, J = 8.4, 5.4 Hz, 1H), 5.39 (d, J = 5.6 Hz, 1H), 4.73 (s, 2H), 4.51 (d, J = 4.2 Hz, 2H), 4.08 (d, J = 13.7 Hz, 1H), 3.87 (t, J = 7.2 Hz, 1H), 3.80 (s, 3H), 3.79 (s, 3H), 3.69 (s, 3H), 3.67 (s, 6H), 3.64 – 3.51 (m, 22H), 3.48 – 3.43 (m, 2H), 3.17 – 2.77 (m, 13H), 2.77 – 2.69 (m, 2H), 2.57 – 2.51 (m, 1H), 2.47 – 2.40 (m, 1H), 2.33 – 2.26 (m, 1H), 2.07 – 1.96 (m, 4H), 1.81 – 1.46 (m, 20H), 0.89 (t, J = 7.4 Hz, 3H). ^13^C NMR (151 MHz, CD_3_OD) δ 175.0, 171.9, 171.1, 169.8, 169.7, 169.5, 159.1, 154.6, 148.9, 143.7, 139.0, 137.9, 137.0, 135.2, 130.9, 130.1, 121.8, 120.7, 117.1, 115.1, 114.4, 113.7, 113.3, 106.6, 77.1, 71.6, 71.5, 71.3, 70.7, 70.4, 68.3, 64.3, 64.2, 61.1, 56.6, 56.5, 53.6, 51.3, 51.2, 51.1, 45.0, 40.9, 40.0, 39.3, 32.3, 29.4, 27.6, 26.3, 25.4, 23.8, 21.9, 12.7. HRMS (ESI) [M+H]^+^ for C_85_H_118_N_8_NaO_18_: 1561.8462, found: 1561.8456.

#### Synthesis of KL-7

**Scheme 10.**
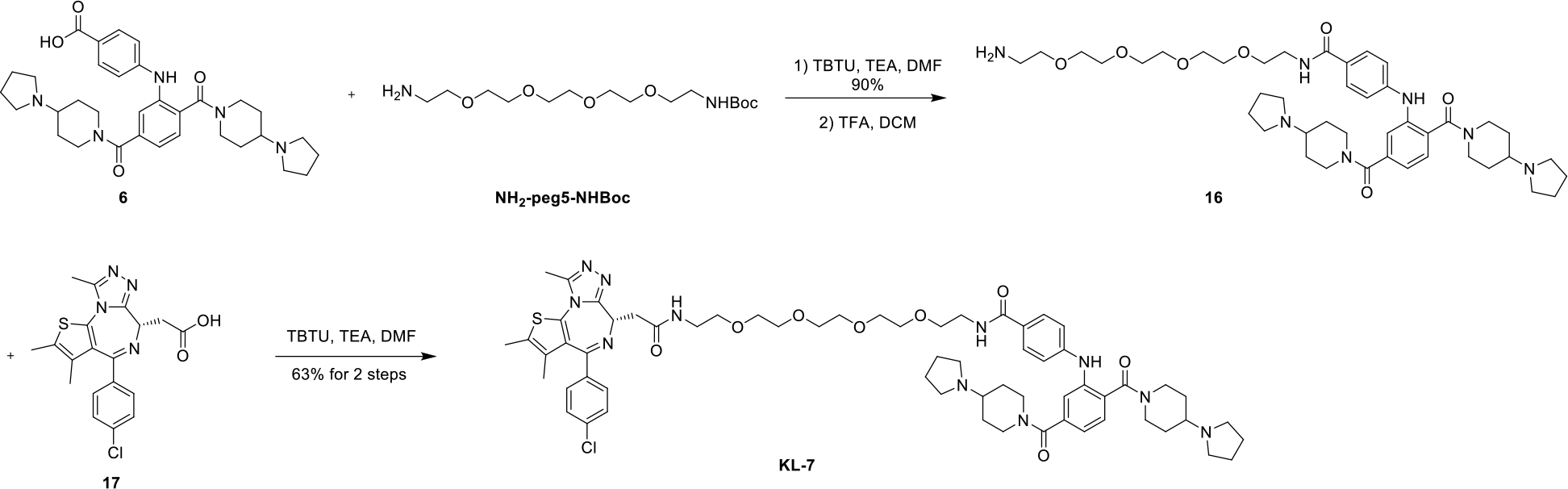
Synthesis of KL-7

**N-[2-[2-[2-[2-(2-aminoethoxy)ethoxy]ethoxy]ethoxy]ethyl]-4-[2,5-bis(4-pyrrolidin-1-ylpiperidine-1-carbonyl)anilino]benzamide (16)**

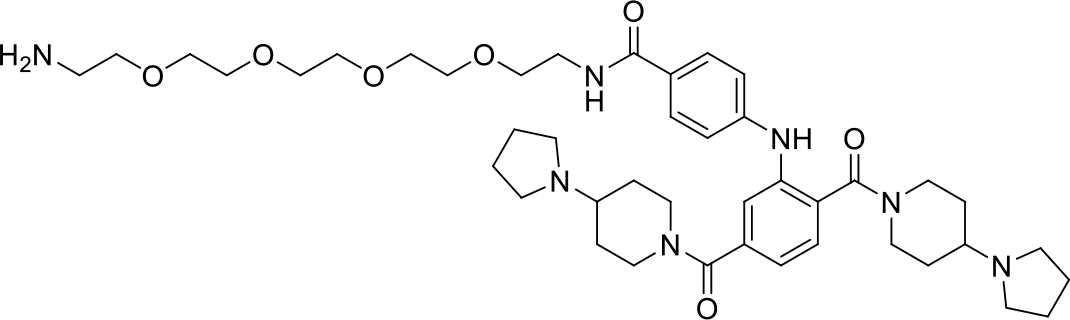

To a mixture of 4-[2,5-bis(4-pyrrolidin-1-ylpiperidine-1-carbonyl)anilino]benzoic acid (**6**) (9.00 mg, 0.0157 mmol) and TBTU (7.56 mg, 0.0235 mmol) in DMF (0.3 mL), a solution of tert-butyl N-[2-[2-[2-[2-(2-aminoethoxy)ethoxy]ethoxy]ethoxy]ethyl]carbamate (5.28 mg, 0.0157 mmol) and TEA (0.0219 mL, 0.157 mmol) in DMF (0.3 mL) was added. The mixture was stirred at rt for 12h. The reaction was quenched by the addition of saturated aq. NaHCO_3_ (2 mL) and extracted with CH_2_Cl_2_ (3 mL x 3). The combined organic extracts were dried over Na_2_SO_4_ and concentrated under reduced pressure. The residue was purified by Prep-TLC (Magic base/CH_2_Cl_2_ = 3:1, magic base = CH_2_Cl_2_/MeOH/NH_3_.H_2_O = 60/10/1) to give tert-butyl N-[2-[2-[2-[2-[2-[[4-[2,5-bis(4-pyrrolidin-1-ylpiperidine-1-carbonyl)anilino]benzoyl] amino]ethoxy]ethoxy]ethoxy]ethoxy]ethyl]carbamate (12.6 mg, 0.0141 mmol, yield: 90.0 %). ^1^H NMR (600 MHz, CD_3_OD) δ 7.77 (d, J = 8.4 Hz, 2H), 7.44 (d, J = 7.8 Hz, 1H), 7.38 (s, 1H), 7.16 (d, J = 7.8 Hz, 1H), 7.10 (d, J = 8.4 Hz, 2H), 4.73 (s, 2H), 4.02 – 3.36 (m, 30H), 3.27 – 2.70 (m, 12H), 2.45 – 1.85 (m, 12H), 1.80 – 1.50 (m, 4H), 1.43 (s, 9H). ^13^C NMR (151 MHz, CD_3_OD) δ 171.2, 169.8, 169.7, 158.4, 147.9, 141.3, 139.0, 130.5, 130.1, 127.4, 121.8, 120.1, 117.5, 80.1, 71.6, 71.6, 71.5, 71.5, 71.3, 71.3, 71.0, 70.7, 70.5, 63.0, 62.3, 52.9, 52.7, 47.0, 41.3, 40.9, 40.2, 31.4, 30.4, 30.2, 29.4, 28.8, 23.9. HRMS (ESI) [M+H]^+^ for C_48_H_74_N_7_O_9_: 892.5548, found: 892.5542.

12.4 mg of the obtained product was dissolved in 1 mL of TFA/CH_2_Cl_2_ (v/v = 1:2). The mixture was stirred at rt for 2h. Concentrated and the crude product (**16**) was used into next step without further purification.

**KL-7**

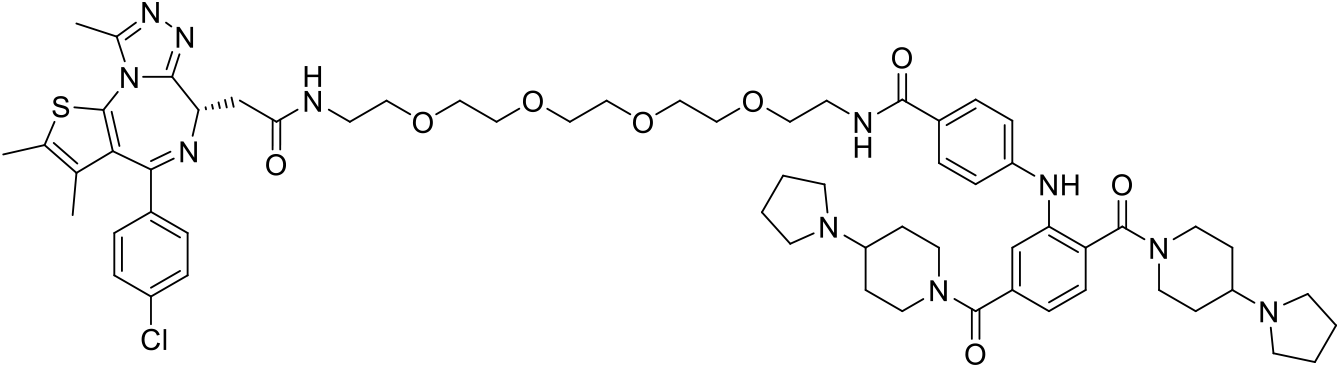

To a mixture of 2-[(9S)-7-(4-chlorophenyl)-4,5,13-trimethyl-3-thia-1,8,11,12-tetrazatricyclo[8.3.0.02,6] trideca-2(6),4,7,10,12-pentaen-9-yl]acetic acid (**17**) (5.57 mg, 0.0139 mmol) and N-[2-[2-[2-[2-(2-aminoethoxy)ethoxy]ethoxy]ethoxy]ethyl]-4-[2,5-bis(4-pyrrolidin-1-ylpiperidine-1-carbonyl)anilino] benzamide (**16**) (11.0 mg, 0.0139 mmol) and TEA (0.0194 mL, 0.139 mmol) in DMF (0.3 mL), TBTU (6.69 mg, 0.0208 mmol) was added. The mixture was stirred at rt for 12h. The reaction was quenched by the addition of saturated aq. NaHCO_3_ (2 mL) and extracted with CH_2_Cl_2_ (3 mL x 3). The combined organic extracts were dried over Na_2_SO_4_ and concentrated under reduced pressure. The residue was purified by Prep-TLC (Magic base/CH_2_Cl_2_ = 3:1, magic base = CH_2_Cl_2_/MeOH/NH_3_.H_2_O = 60/10/1) to give **KL-7** (10.3 mg, 0.00877 mmol, yield: 63.1 %). ^1^H NMR (600 MHz, DMSO) δ 10.68 (s, 1H), 8.28 (t, J = 5.6 Hz, 2H), 8.20 (s, 1H), 7.74 (d, J = 8.3 Hz, 2H), 7.48 (d, J = 8.3 Hz, 2H), 7.42 (d, J = 8.3 Hz, 2H), 7.33 (d, J = 7.8 Hz, 1H), 7.23 (s, 1H), 7.10 – 6.96 (m, 3H), 4.53 – 4.33 (m, 3H), 3.80 – 3.42 (m, 20H), 3.31 – 3.16 (m, 6H), 3.12 – 2.66 (m, 9H), 2.59 (s, 3H), 2.40 (s, 3H), 2.12 – 1.68 (m, 12H), 1.62 (s, 3H), 1.58 – 1.32 (m, 4H). ^13^C NMR (151 MHz, DMSO) δ 170.1, 168.5, 167.1, 166.2, 163.4, 155.5, 150.3, 146.7, 139.5, 137.9, 137.2, 135.7, 132.7, 131.1, 130.6, 130.3, 130.0, 129.3, 129.1, 128.9, 126.1, 120.7, 118.8, 116.1, 70.2, 70.2, 70.1, 70.0, 69.6, 69.5, 60.9, 60.7, 54.3, 50.9, 50.8, 45.8, 45.2, 39.1, 37.9, 28.3, 23.2, 23.2, 14.5, 13.1, 11.7. HRMS (ESI) [M+H]^+^ for C_62_H_81_ClN_11_O_8_S: 1174.5679, found: 1174.5673.

## Supporting Figures

**Figure S1.**
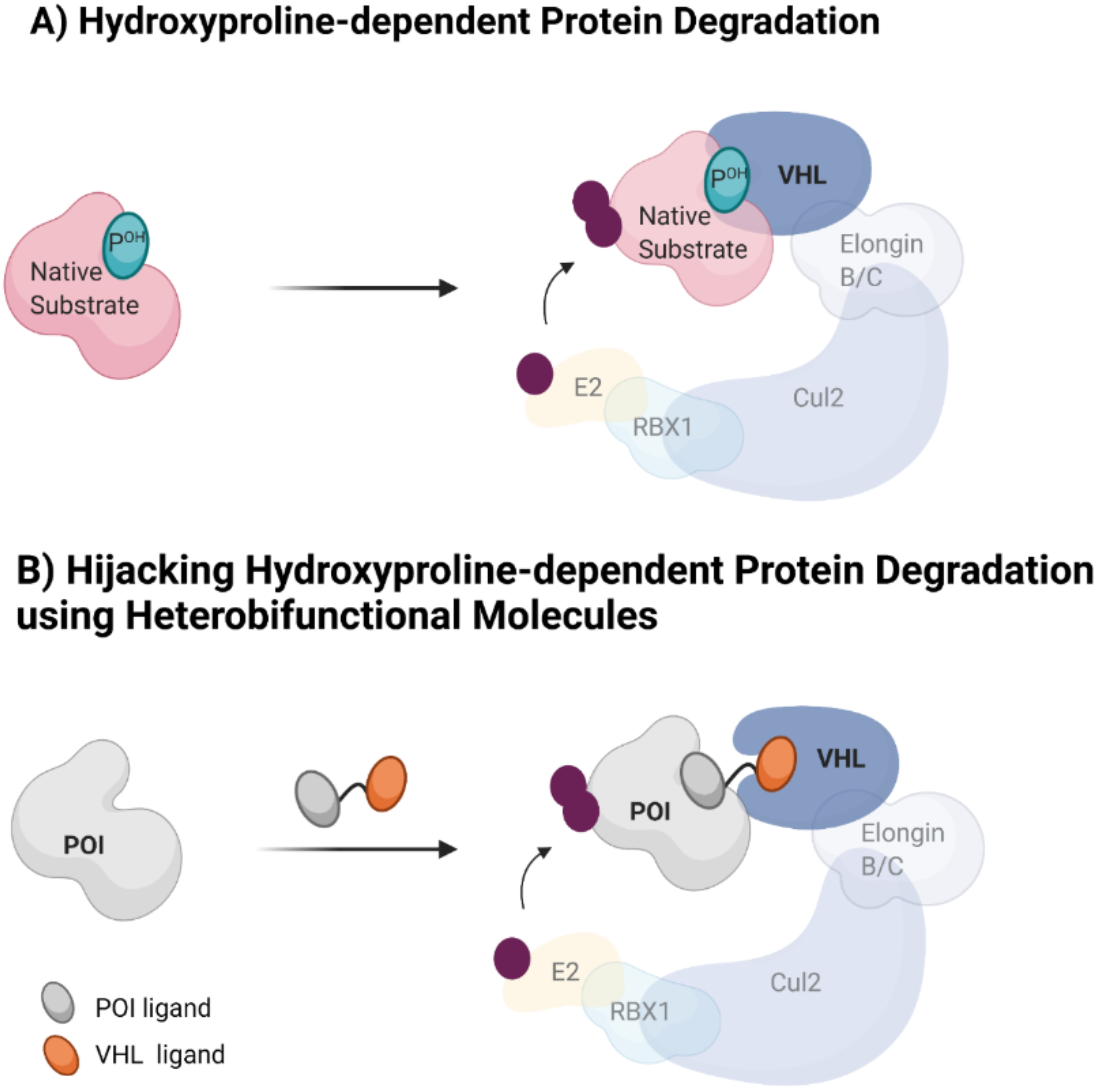
Hijacking hydroxy-proline binding to VHL using chimeric molecules. A) Natural hydroxyproline-dependent protein degradation system. VHL binds to prolyl-hydroxylated proteins such as HIF-1α and targets them for proteasomal degradation via the Cul2 E3 ligase complex. B) Hijacking hydroxyproline-dependent protein degradation using heterobifunctional molecules. VHL-recruiting PROTACs induce ternary complex formation between protein of interest (POI) and VHL E3 ligase, thus promoting ubiquitination and degradation of the POI.

**Figure S2.**
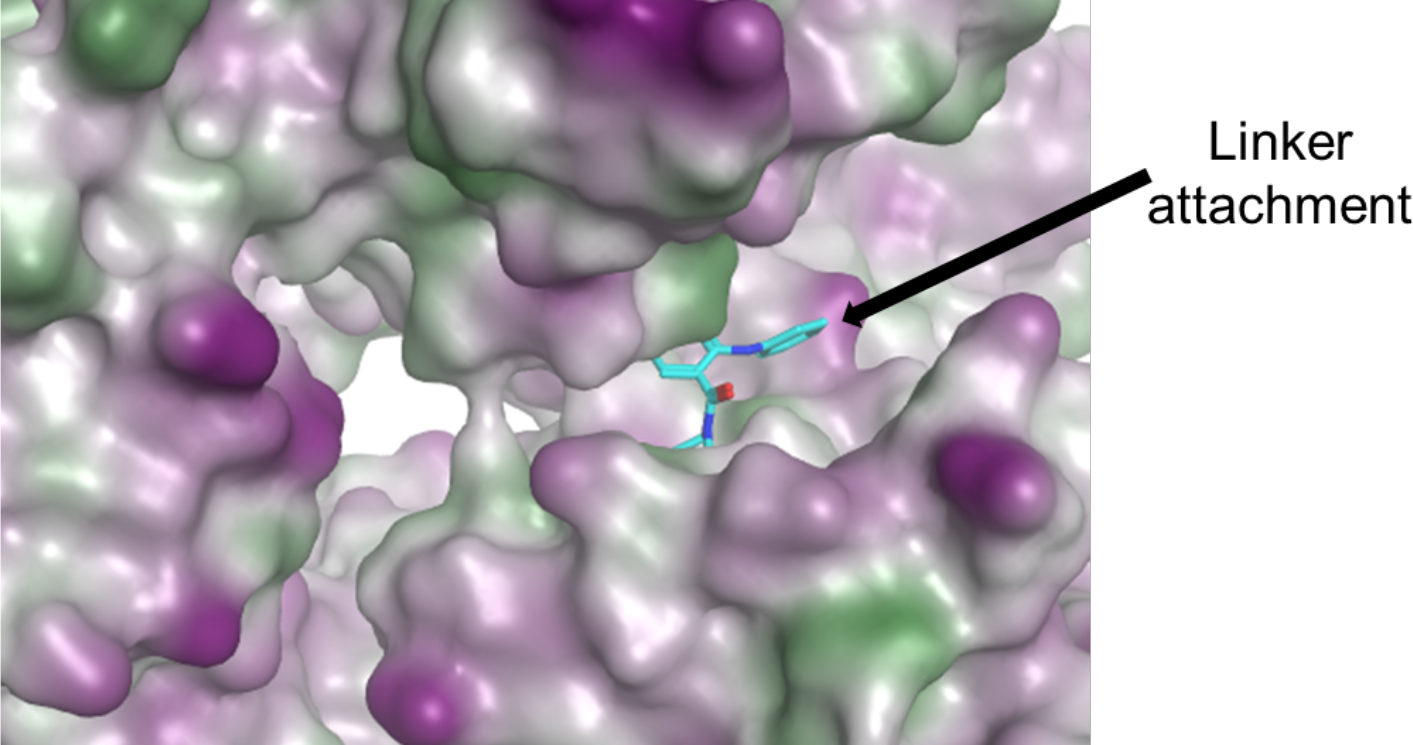
Crystal structure of UNC1215 bound to L3MBTL3 (PDB: 4FL6). UNC1215 is shown in cyan, the black arrow indicates the point of linker attachment.

**Figure S3.**
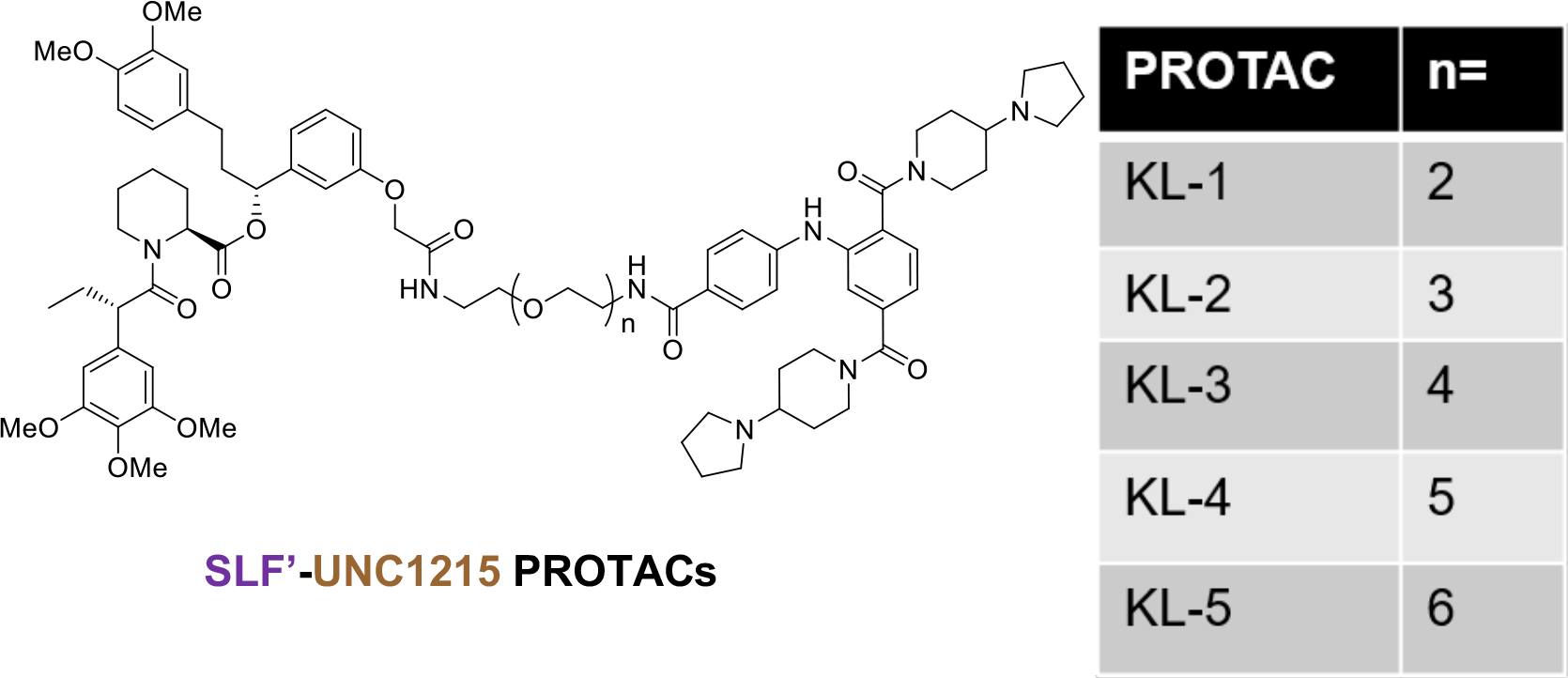
Chemical structure of SLF’-UNC1215 PROTACs. FKBP12^F36V^ selective bumped ligand (SLF’) is fused to the L3MBTL3 ligand (UNC1215) via varying linker lengths, n= 2-6.

**Figure S4.**
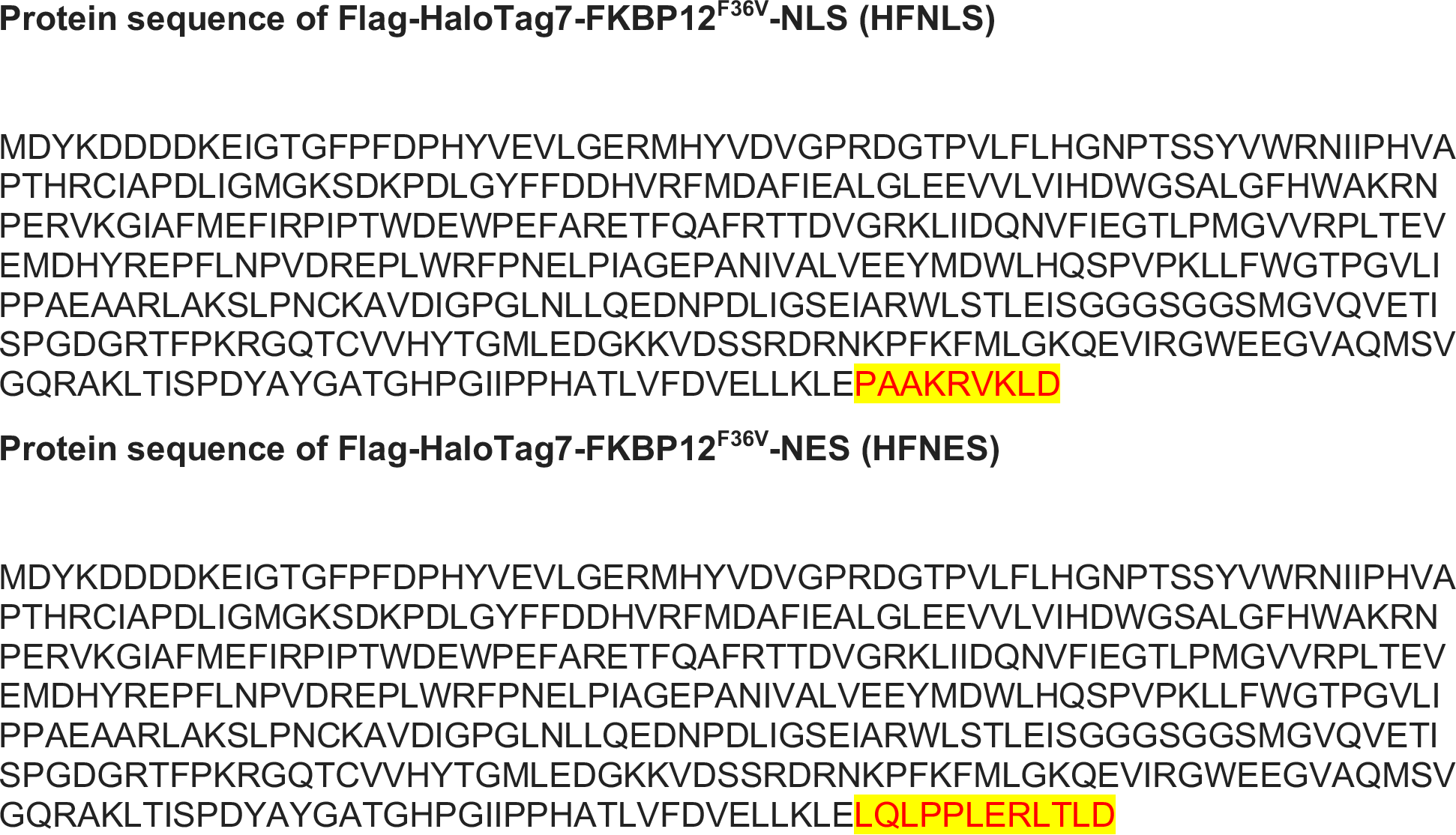
Protein sequences of HFNLS (FLAG-HaloTag7-FKBP12^F36V^-NLS) and HFNES (FLAG-HaloTag7-FKBP12^F36V^-NES). NLS and NES sequence of each protein is highlighted in yellow.

**Figure S5.**
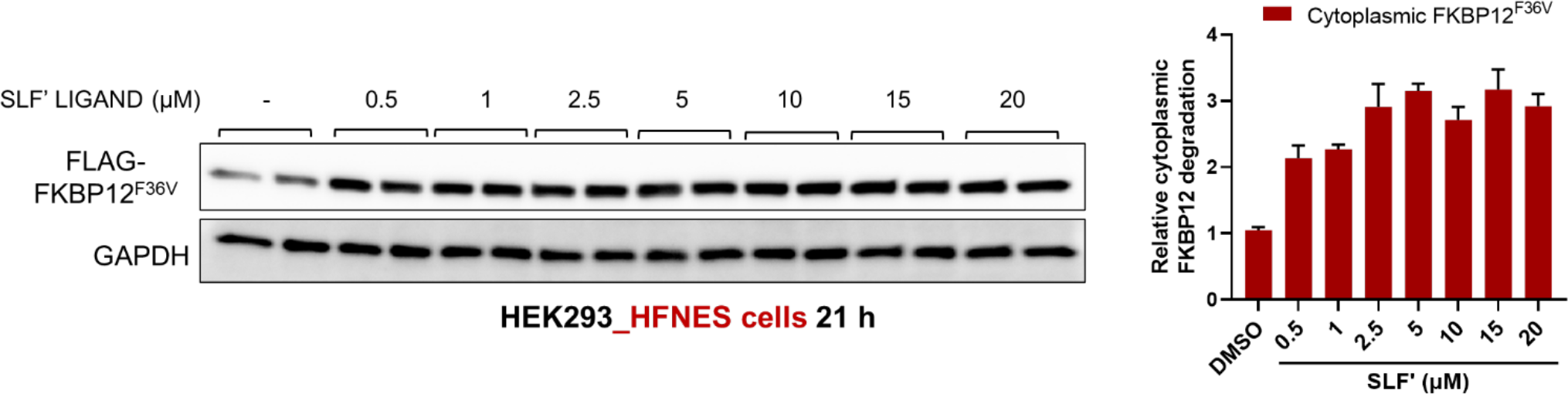
Dose response of SLF’ in HFNES cells. SLF’ ligand induces stabilization of cytoplasmic FKBP12 in HFNES cells. HEK293_HFNES cells were treated with increasing concentration of SLF’ for 21h and then analyzed by blotting with FLAG and GAPDH antibodies. Quantitation is on the right and the quantified data represent mean ± s.e.m, n = 2.

**Figure S6.**
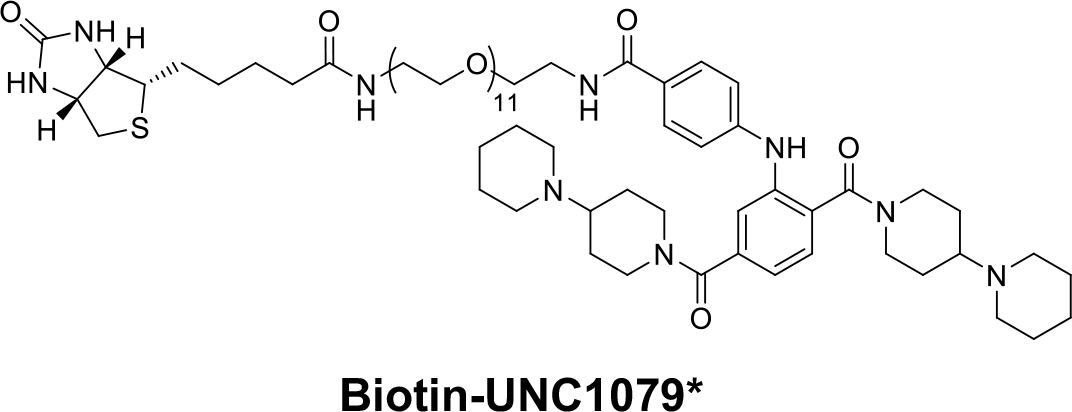
Chemical structure of Biotin-UNC1079*.

**Figure S7.**
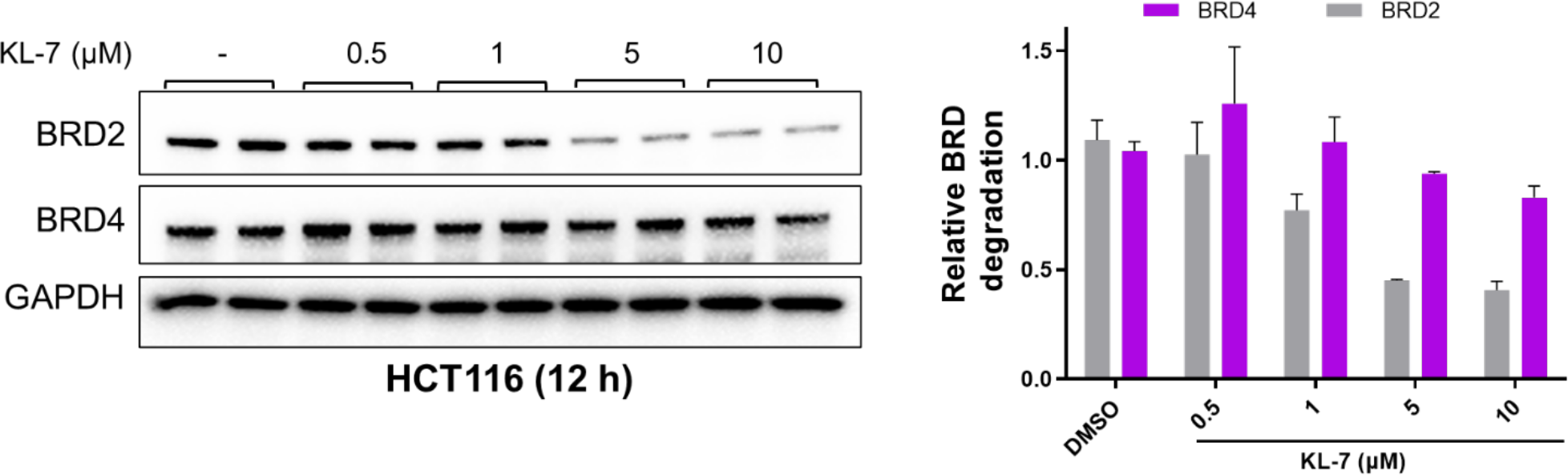
Dose response of KL-7 in HCT116 cells. KL-7 promotes selective BRD2 degradation over BRD4 in HCT116 cells. HCT116 cells were treated with increasing concentrations of KL-7 for 12 h. Quantitation is on the right and the quantified data represent mean ± s.e.m (n = 2).

**Figure S8.**
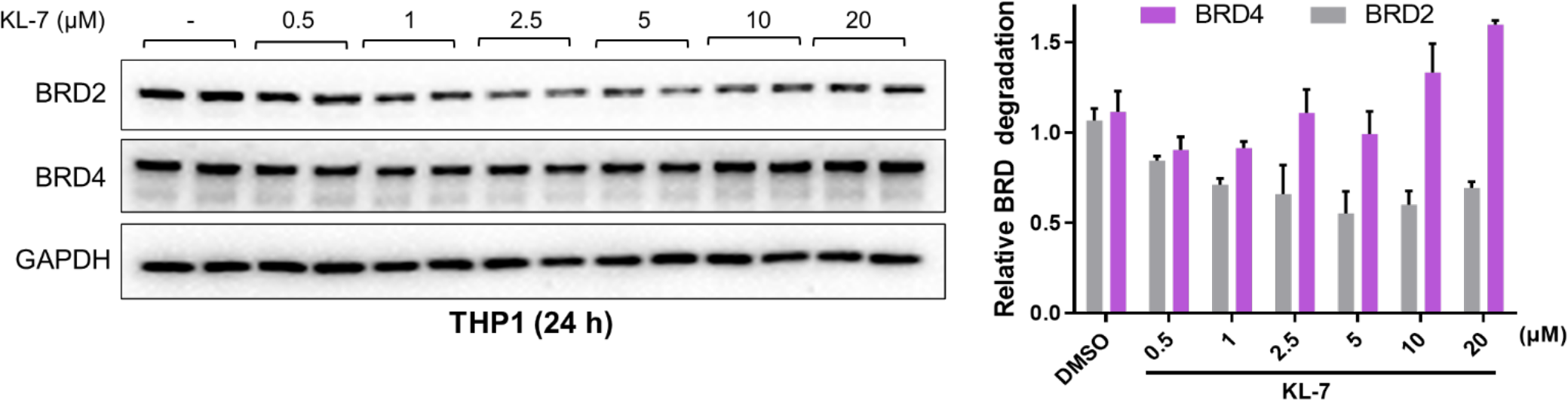
Dose response of KL-7 in THP1 cells. KL-7 promotes selective BRD2 degradation over BRD4 in THP1 cells. THP1 cells were treated with increasing concentrations of KL-7 for 24h. Quantitation is on the right and the quantified data represent mean ± s.e.m (n = 2).

**Figure S9.**
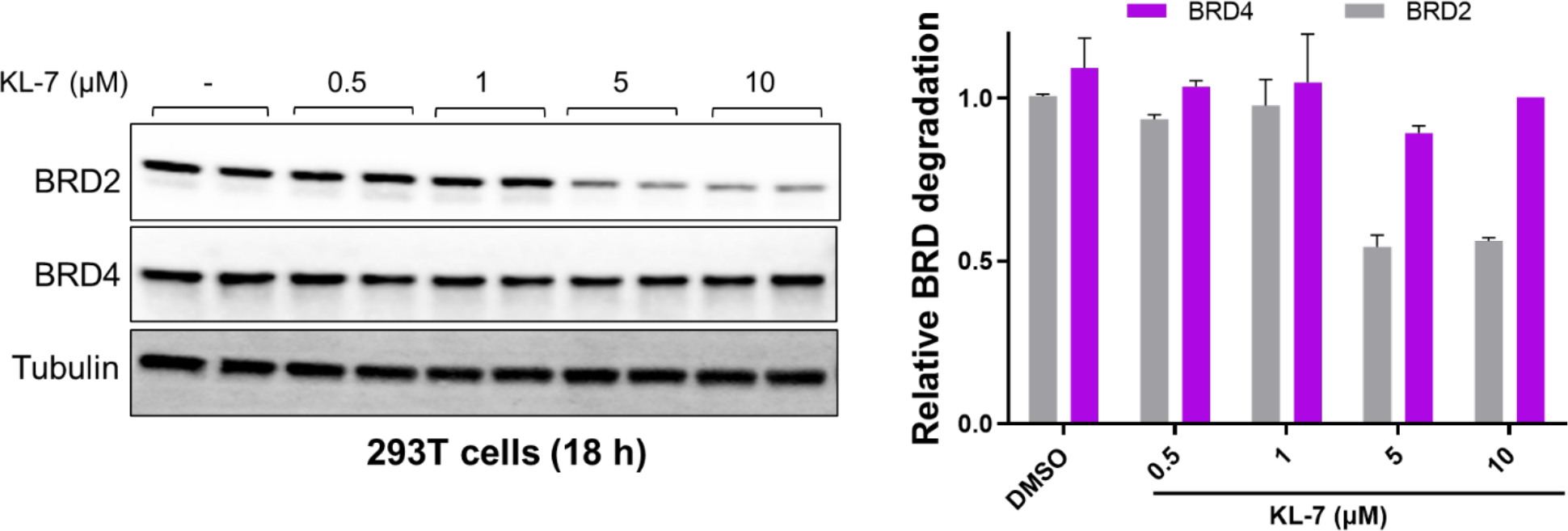
Dose response of KL-7 in HEK293T cells. KL-7 promotes selective BRD2 degradation over BRD4 in HEK293T cells. HEK293T cells were treated with increasing concentrations of KL-7 for 18 h. Quantitation is on the right and the quantified data represent mean ± s.e.m (n = 1).

